# Microglial adipose triglyceride lipase regulates neuroinflammatory and behavioural responses to LPS

**DOI:** 10.1101/2023.11.30.569441

**Authors:** Josephine Louise Robb, Arturo Israel Machuca-Parra, Frédérick Boisjoly, Adeline Coursan, Romane Manceau, Danie Majeur, Demetra Rodaros, Khalil Bouyakdan, Karine Greffard, Jean-François Bilodeau, Anik Forest, Caroline Daneault, Matthieu Ruiz, Cyril Laurent, Nathalie Arbour, Sophie Layé, Xavier Fioramonti, Charlotte Madore-Delpech, Stephanie Fulton, Thierry Alquier, Representative of consortium

## Abstract

Adipose triglyceride lipase (ATGL), the enzyme that catalyses the rate-limiting step of triglyceride lipolysis, regulates inflammation in peripheral tissues. ATGL has been associated with both pro- and anti-inflammatory responses in different tissues suggesting its actions are dependent on cell type. Recent studies in microglia and macrophages suggest that lipid droplets (LD), a triglyceride storing organelle, and LD lipolysis via ATGL are important components of inflammatory responses. Here, we determined the impact of ATGL inhibition and microglia-specific ATGL loss-of-function on inflammatory and behavioural responses to acute pro-inflammatory insult. First, we evaluated the impact of lipolysis inhibition on lipopolysaccharide (LPS)-induced expression and secretion of cytokines in mouse primary microglia cultures. LPS led to LD accumulation in microglia and altered the expression of lipolysis regulators. The pan-lipase inhibitor ORlistat alleviated LPS-induced expression of IL-1β and IL-6. Specific inhibition of ATGL by ATGListatin had similar anti-inflammatory action on cytokines expression and secretion in both neonatal and adult microglia cultures. Second, targeted and untargeted lipidomic studies revealed that ATGL inhibition reduced LPS-induced generation of pro-inflammatory prostanoids and affected ceramide profile. Finally, the role of ATGL in neuroinflammation was assessed in a novel mouse model with inducible ATGL deletion specifically in microglia. Loss of microglial ATGL in adult male mice dampened LPS-induced expression of IL-6 and reduced LPS-induced sickness behaviour. Together, our results demonstrate that pharmacological inhibition or loss of ATGL-mediated triglyceride lipolysis reduces LPS-induced inflammation to suggest that inhibition of lipolysis plays a beneficial role in neuroinflammation.

## 1 Introduction

Lipid droplets (LDs) are dynamic organelles primarily recognised as neutral lipid storage sites for cholesterol esters and triglycerides (TG). TG hydrolysis releases fatty acids for use as energy substrates. LDs are also a source of fatty acids that are precursors for the synthesis of lipid-mediators of inflammation including eicosanoids (Jarc & Petan, 2020). The phospholipid monolayer surrounding LDs contains glycerolipid lipases, which catalyse TG hydrolysis, and regulatory proteins such as perilipins (PLIN) (Olzmann & Carvalho, 2019). The first committed step of TG hydrolysis is catalysed by adipose triglyceride lipase (ATGL). ATGL hydrolyses TG (when interacting with the enhancer ABHD5 (CGI-58)) to generate a diacylglyceride (DAG) and a free fatty acid (Eichmann et al., 2012; Eichmann & Lass, 2015; Zechner et al., 2009). Subsequent steps of DAG and monoacylglyceride (MAG) hydrolysis are catalysed by the hormone-sensitive lipase (HSL) and MAG lipase (MAGL) respectively (Jarc & Petan, 2020). It is now well established that LDs and neutral glycerolipid lipases have a broad role in cell metabolism, signalling and physiology (Jarc & Petan, 2020; Olzmann & Carvalho, 2019).

Recent studies have reported LD accumulation in microglia, the resident immune cells of the brain, in response to inflammatory insults, ageing and neurodegeneration (Khatchadourian et al., 2012; Li et al., 2023; Loving et al., 2021; Marschallinger et al., 2020; Qin et al., 2023), thereby suggesting a strong association between LD accumulation in microglia and neuroinflammation. *In vitro,* accumulation of LDs in microglia in response to lipopolysaccharide (LPS) is currently thought to be caused by the recruitment of perilipin 2 (PLIN2) to the LD membrane and subsequent PLIN2-dependent inhibition of ATGL activity (Khatchadourian et al., 2012; Wang et al., 2011). This suggests that ATGL activity is regulated by inflammatory signals and contributes to LD accumulation in microglia.

In peripheral tissues and cells, inhibition or loss of ATGL activity can promote inflammation in a cell type-dependent manner. For example, ATGL genetic loss-of-function in white or brown adipocytes increases interleukin (IL)-6 and tumour necrosis factor (TNF)-α secretion (Kotzbeck et al., 2018; Lettieri Barbato, Aquilano, et al., 2014; Lettieri Barbato, Tatulli, et al., 2014). Similarly, loss of ATGL enhances the expression of inflammatory markers in the liver such as macrophage chemoattractant protein (MCP)-1, TNF-α and IL-1β in response to LPS (Jha et al., 2014). In contrast, ATGL deletion in macrophages leads to a decrease in IL-6 signalling and an increase in IL-10 and transforming growth factor (TGF)-β in response to MCP-1 stimulation or in atherosclerotic mice suggesting that loss of ATGL activity promotes anti-inflammatory actions (Aflaki et al., 2011; Lammers et al., 2011). On the contrary, stimulation of TG lipolysis in macrophages through activation of ATGL induced by the loss of the inhibitor HILPDA increases prostaglandin E_2_ (PGE_2_), TNF-α and IL-6 production (van Dierendonck et al., 2022). Recently, it was shown that inhibition of ATGL reduces pro-inflammatory responses in primary microglia to suggest microglial ATGL may play a broad role in microglia reactivity. Despite these findings, the contribution of microglial ATGL to neuroinflammation and associated behavioural deficits in response to pro-inflammatory insults as well as the underlying mechanisms remain unknown. Using complementary tools and models, we demonstrate here that pharmacological inhibition of ATGL in primary microglia or specific microglial ATGL loss-of-function in the mouse brain robustly reduces microglial pro-inflammatory and behavioural responses to LPS.

## 2 Methods

### 2.1 Animal ethics

All animal studies were conducted in accordance with guidelines of the Canadian Council on Animal Care (protocol #CM19018TAs). Adult microglia cultures were conducted in accordance with the European directive 2010/63/UE and approved by the French Ministry of Research and local ethics committees (APAFIS#: 33951). C57BL6/J mice purchased from Jackson Laboratories were used to generate pups for the cultures of primary microglia from the forebrain. Mice were group housed on a 12 h dark-light cycle at 22-23 °C, with *ad libitum* access to standard irradiated chow diet (Teklad) and water. For studies in adult transgenic mice, age-matched littermates were used and individually housed in a reverse light cycle after genotyping. Only treatment-naive mice were used at the time of study. ATGL^fl/fl^ mice were kindly donated by Dr Grant Mitchell and were backcrossed at least 6 generations on the C57BL/6J genetic background (C57BL/6J, 000664). Female ATGL^fl/fl^ mice on the C57BL/6J background were bred with male mice expressing tamoxifen-inducible CreERT2 recombinase and YFP under the control of the mouse CX3CR1 promoter [B6.129P2(Cg)-Cx3cr1tm2.1(cre/ERT2)Litt/WganJ, 021160], obtained from The Jackson Laboratory (Littman, 2013; Sahasrabuddhe & Singhee Ghosh, 2022). ATGL^+/+^;CX3CR1-CreERT2 (hemizygous) controls (CX3CR1^ATGL^ WT) and ATGL^fl/fl^;CX3CR1-CreERT2 (CX3CR1^ATGL^ KO) were obtained by breeding female ATGL^fl/+^ with male ATGL^fl/+^;CX3CR1-CreERT2 (hemizygous) to obtain littermates of all genotypes.

#### 2.1.1 Tamoxifen delivery

Tamoxifen (Sigma) was dissolved in 10% ethanol, 90% corn oil by heating at 50 °C. Two doses of 10 mg tamoxifen were delivered by oral gavage at 48 h intervals. All experimental mice (CX3CR1^ATGL^ WT and CX3CR1^ATGL^ KO) were injected with tamoxifen. Experiments began 4 weeks post-tamoxifen to allow repopulation of non-recombined macrophages in the periphery to ensure a microglia-specific knockout (Parkhurst et al., 2013).

#### 2.1.2 Validation of the microglial ATGL KO mouse model

Validation was performed using FACS sorting. Digestion, FACS and analysis were carried out as described by Legroux *et al*. (2015). Briefly, brains and spleens from Cx3CR1^ATGL^ WT or KO mice were digested with Collagenase D (Roche) and DNase I. Cells were recovered and separated in CD11b^+^ microglial/macrophages or CD11b^-^ non-microglial fractions using Percoll^TM^ gradient followed by the pull-down cell selection method, EasyStep Mouse CD11b positive Selection Kit II (StemCell Technologies, 18970A), as per the manufacturer’s instructions. Surface staining for CD45 PE-Cy7 (clone 30-F11; #552848, BD Biosciences) was carried out as previously described (Legroux et al., 2015). Cell viability was determined using Live/Dead^TM^ (L34957, Invitrogen). The following fractions were sorted by flow cytometry: CD45^+^/YFP^+^ gated events from the CD11b^+^ brain fraction represent microglial cells (CD11b^+^/YFP^+^); CD45^-^/YFP^-^ gated events from the CD11b^-^ fraction represent non-microglial neural cells (CD11b^-^/YFP^-^); CD45^+^/YFP^-^ gated events from the CD11b^+^ fraction from the spleen represent macrophages cells (CD11b^+^/YFP^-^). RNA was extracted from the resulting fractions with the RNeasy Plus Micro Kit (74034; Qiagen), as per manufacturer’s instructions, and gene expression was quantified and characterised by qPCR as described in section 2.5.

#### 2.1.3 Behavioural testing

Animals were administered saline or 0.83 mg/kg LPS *i.p.* 12 h prior to behavioural testing. Behavioural testing was performed during the dark cycle. Emotionality scores were calculated as described by Guilloux *et al*. (2011). Briefly, z scores for each test were calculated as follows:

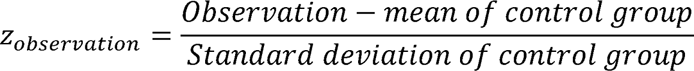

These Z_observation_ scores were combined for the elevated plus maze (EPM) and light-dark box (LDB) tests:

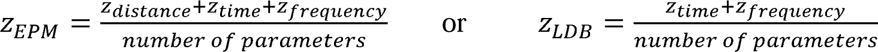

These resulting numbers were combined into an emotionality score:

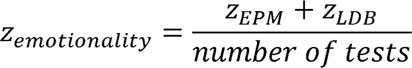

##### 2.1.3.1 Elevated plus maze

Test was performed as previously reported (Sharma & Fulton, 2013). Briefly, animals were placed in the centre of the maze facing the open arm away from the experimenter and movement was tracked using and overhead camera for 5 minutes. Place preference and movement was analysed using Ethovision as previously described (Sharma & Fulton, 2013).

##### 2.1.3.2 Light-Dark box

This test was performed in a chamber comprising of two sections: an enclosed dark zone of opaque plexiglass, and a light zone of clear plexiglass and open at the top. Both sections were 13.7 cm × 13.7 cm × 20.3 cm. An opening at the base of the dark section allowed free movement between sections. Animals were placed in the centre of the box adjacent to the entry to the dark zone. Movement was tracked using and overhead camera for 5 minutes. Place preference and movement was analysed using Ethovision.

### 2.2 Cell culture

#### 2.2.1 Primary neonatal mouse microglia

Primary neonatal mouse microglia were isolated from the forebrain of P1-2 mice. Cerebellum, olfactory bulbs, and meninges were removed before rough homogenization with a blade. The homogenate was centrifuged at 500 g for 1 minute and supernatant discarded. Homogenate was digested for 15 minutes in enzymatic solution (0.2 µm filter sterilised 4.5 U/mL papain [Worthington Biochemicals], 1× DNAse [Worthington Biochemicals], in PBS) at 37 °C. The digest was centrifuged at 500 g for 2 minutes and supernatant discarded. The digest was mechanically disrupted with a serological pipette in complete media (DMEM [Gibco; 11965-092] 4.5 g/dL glucose, 10% FBS [Gibco], 50 U/mL penicillin, 50 µg/mL streptomycin [Gibco]). Cell suspension was filtered through a 70 µm cell strainer, centrifuged at 500 g for 5 minutes and resuspended in complete media. Cell suspension was plated in T75 flasks (50% forebrain/flask) and allowed to adhere for 7 days. Media was changed when astrocyte monolayer reached confluency. When microglial cells began to detach from the astrocyte layer after another 5 days in culture, microglia were dislodged by gently tapping the side of the flask and media containing microglia was centrifuged at 500 g for 10 minutes at RTP. Cells were counted using a hemocytometer and seeded at an appropriate density. Complete media was added to the experimental dishes and microglia were allowed to adhere for 24 h prior to the start of the experiment.

#### 2.2.2 Primary adult mouse microglia

For adult microglia primary cultures, male C57BL/6J mice (12-15 weeks old) were euthanized by intraperitoneal pentobarbital injection (Exagon®, 300 mg/kg), 30 minutes post-buprenorphine administration (0.1 mg/kg, subcutaneously). Microglia isolation was performed as previously described (Herron et al., 2022). In brief, mice were transcardially perfused with ice-cold Hanks’ Balanced Salt Solution (HBSS, Gibco), followed by mechanical dissociation of forebrain tissue. Single cell suspensions were prepared and centrifuged over a 37%/70% discontinuous Percoll gradient (GE Healthcare), mononuclear cells were isolated from the interface. Isolated adult microglia were cultured using a modified version of the protocol described by Butovsky *et al*., with a physiological level of glucose in the culture medium (Butovsky et al., 2014). Microglia were cultured for 5 days at 37 °C, 5% CO2 in poly-D-Ornithine coated 48-well plates (9-10×10^4^ cells/well in 0.5 mL) and grown in supplemented DMEM SILAC medium (Gibco), containing Glucose 2.5 mM, Lysine 0.5 mM, Arginine 0.7 mM, Glutamine 2.5 mM, mouse recombinant carrier free MCSF 10 ng/mL (R&D Systems) and human recombinant TGF-β1 50 ng/mL (Miltenyi Biotec).

#### 2.2.3 Cell culture treatments

For neonatal cultures, culture media was replaced 2 h prior to treatment with pre-treatment medium (DMEM [Winsent; 319-061-CL] containing 5 mM glucose). At time of treatment, pre-treatment medium was replaced with DMEM (5 mM glucose) containing Orlistat (50 µM; MedChemExpress), ATGListatin (50 µM; MedChemExpress), etomoxir (10 µM, Caymen Chemicals) or vehicle (DMSO 0.1% v/v) with or without LPS (0.1 µg/mL; O127:B8; Sigma). For adult microglial cultures, microglia were pre-conditioned during 24 h without TGF-β1 and MCSF. Cells were treated on Day 6 with 0.1 µg/mL LPS. At the same time, ATGListatin (50 µM) or vehicle (DMSO 0.1% v/v) was added for 6 h.

### 2.3 Lipid droplet imaging

Primary microglia derived from pups were plated at a density of 1×10^5^ on 13 mm diameter glass coverslips in 24 well plates. After 6 h treatment, cells were fixed for 15 minutes with 4% paraformaldehyde at 37 °C. Intracellular LDs were visualised using the neutral lipid stain BODIPY 493/503 (10 µM; DMSO; Invitrogen). Hoescht 33342 (6 µg/mL; Invitrogen) was used to identify nuclei. After staining, cells were mounted in ProLong Gold antifade mountant (Invitrogen) and allowed to set prior to imaging (64x oil-immersion lens; Leica SP5 multiphoton system). LDs were manually counted in ImageJ (FIJI; (Schindelin et al., 2012; Schneider et al., 2012)).

### 2.4 Immunoblotting

Primary mouse microglia were plated at a density of 1×10^6^ in T25 flasks. After treatment, cells were lysed using modified RIPA buffer and centrifugation at 21100 g at 4 °C for 20 minutes. Protein content was estimated using the Bradford method microassay procedure using a BSA standard curve (BioRad Protein Assay), as recommended by the supplier. Protein concentrations were standardised with modified RIPA buffer and lamaelli buffer. Protein separation of 10 µg sample was carried out using the BioRad mini-protean system using 12.5% vol/vol gels and SDS-PAGE. Precision Plus Dual Colour standard (BioRad) was used to estimate protein size. Protein was transferred onto 0.2 µm nitrocellulose. Membranes were blocked 1 h with 5% wt/vol milk in TBS and probed overnight with primary antibody (anti-ATGL: 2138S, 1:1000; anti-GAPDH: 2118S, 1:10000 [CST]). TBS-Tween 0.1% vol/vol (TBS-T) was used to remove unbound antibody between each binding steps. Membranes were incubated with secondary antibody goat anti-rabbit conjugated to HRP (1:5000, 5% wt/vol milk in TBS-T; #170-6515, BioRad) for 1 h. Protein was detected by incubation with SuperSignal West Pico Plus Chemiluminescence substrate (Invitrogen) for 5 minutes prior to imaging for 5 minutes with ChemiDoc Imaging system. For reblotting, membranes were stripped using Restore stripping buffer for 15 minutes prior to reblocking.

### 2.5 RNA analyses

#### 2.5.1 Cell and tissue collection for RNA extraction

Primary mouse microglia were plated at a density of 3×10^5^ in 6 well plates and treated for 6 h prior to RNA extraction with Trizol (Invitrogen; Taïb et al., 2013). Adult microglia were treated as described in section 2.2.3 prior to extraction of total RNA using RNeasy Plus Mini Kit (Qiagen, 74106) according to the manufacturer’s protocol.

CX3CR1^ATGL^ WT or KO mice were administered with either a single 0.83 mg/kg or two 0.5 mg/kg injections of LPS *i.p.* (O127:B8) at 24 h intervals, or 0.9% saline vehicle (1 mL/kg, volume matched to LPS). Animals were euthanized by decapitation under isoflurane anaesthesia 3 h after the final injection for tissue collection. Microdissections of brain regions were taken from the cortex, hippocampus and mediobasal hypothalamus (MBH) for RNA extraction. RNA was extracted from tissue samples using the Trizol method (Invitrogen; Taïb et al., 2013).

#### 2.5.2 RNA extraction, cDNA synthesis and RT-qPCR

Extracted RNA was checked for purity and concentration using the Nanodrop 2000. For RNA extracted from neonatal microglia or tissue, cDNA was synthesised from 1 µg of RNA with M-MuLV reverse transcriptase (Invitrogen) and random hexamers as specified by the manufacturer. cDNA was diluted 1:10 prior to qPCR. qPCR master mix was composed of QuantiFast SYBR Green PCR kit (Qiagen) and forward and reverse primers (1 µM) for the relevant gene of interest. Primer sequences used are provided in Table 1. qPCR was performed for 40 cycles using the Corbett Rotor-Gene 6000 (Qiagen) and quantified using the standard curve method using Rotorgene Q series software (v2.3.1). Genes of interest were normalised to either 18S or β-actin (*in vitro*), or the geometric mean of β-actin and cyclophilin (*in vivo*) as was determined by the stability of the respective housekeeping genes using NormFinder (Andersen et al., 2004). Gene expression is represented as fold change from mean of LPS-treated control groups after normalisation.

**Table 1.**
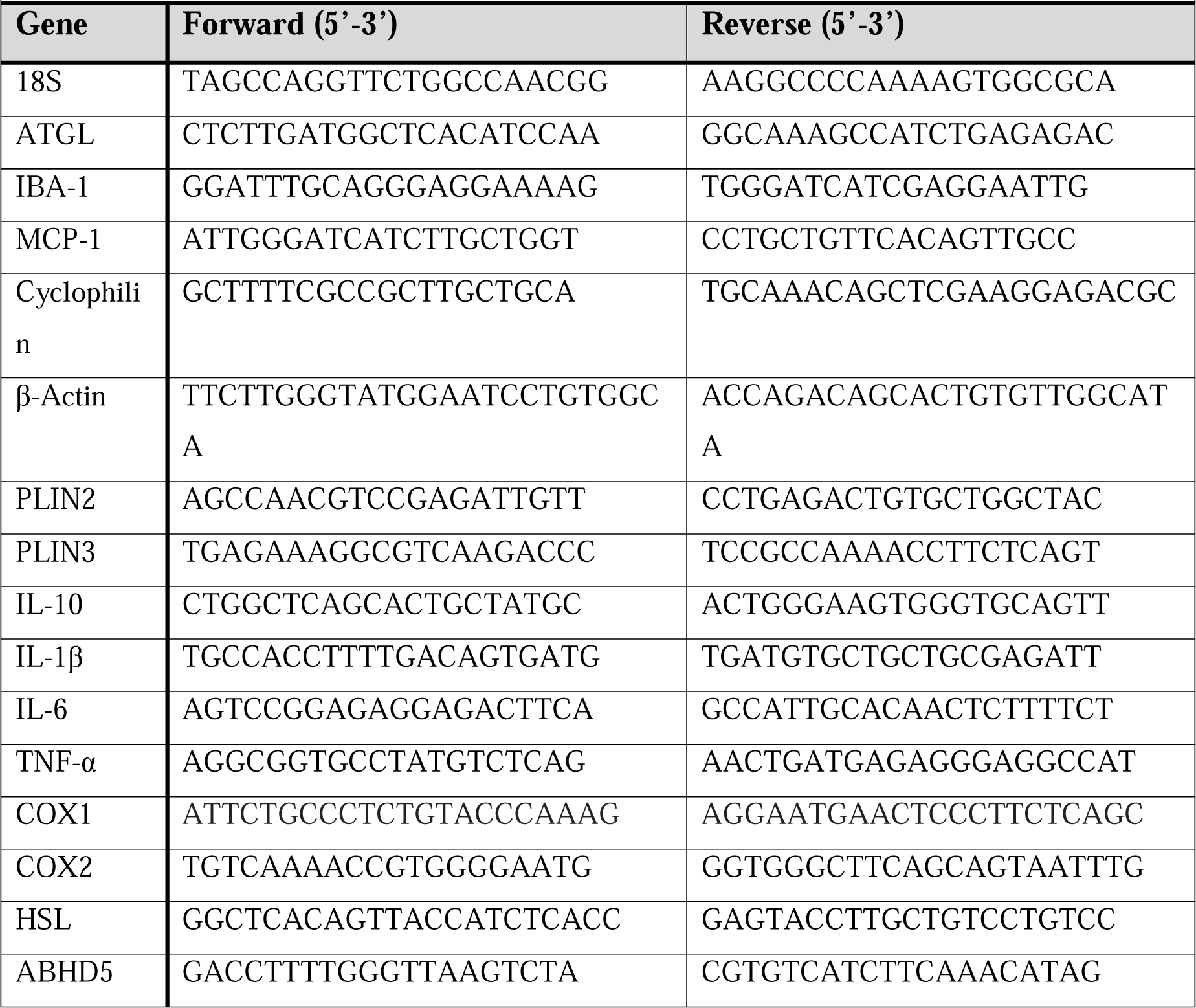
*Primers* Primer sequences used for SYBRGreen qPCR.

For adult microglia, 10µL of RNA was reverse transcribed using Superscript IV VILO (Invitrogen, Life Technologies, France). cDNA was amplified using FAM-labeled Taqman® probes (Applied Biosystems/Thermo Fisher Scientific), [ccl2 (Mm00441242_m1), il-10 (Mm01288386_m1), il-1b (Mm00434228_m1), il-6 (Mm00446190_m1), tnf-a (Mm00443258_m1), B2m (Mm00437762_m1) used as reference gene]. Real-time PCR reaction was performed in duplicate using a LightCycler® 480 instrument II (Roche). Data are normalized relative to B2m (2^(−ΔCt)^).

### 2.6 Cytokine secretion

Primary microglia derived from pups were seeded at a density of 4×10^4^ in 96 well plates and treated for 3-24 h prior to collection of the extracellular media. MCP-1 and IL-6 secretion into the extracellular media was quantified using DuoSet ELISAs (Bio-techne; DY479-05 and DY406-05 respectively), as directed by the manufacturer.

### 2.7 Endogenous oleate oxidation assay

Microglial cells (1×10^6^) derived from pups were seeded in T25 flasks. Cells were incubated in phenol red free DMEM (10 mM glucose) containing 0.25 mM oleate (Sigma) pre-complexed to 0.3% BSA and 0.1 μCi/ml [1-^14^C] oleate (Perkin Elmer) for 24 h to label all cellular lipids including TG as described previously (Taïb et al., 2013). Prior to treatment, exogenous oleate was removed by 2 washes with PBS. Cells were then treated with β-oxidation media (phenol red free DMEM containing 1 mM carnitine, 0.3% BSA, 2.5 mM glucose) containing vehicle, ATGListatin (50µM) or 10 µM etomoxir and incubated for 3 h.

### 2.8 Targeted lipidomics

#### 2.8.1 Materials

Leukotriene(LT) B4, lipoxin(LX) A4, 6(S)-LXA4, 15(R)-LXA4, LXB4, maresin-1 (MaR1), protectin 1 (PD1), PDx, Resolvin D1 (RvD1), 17(R)-RvD1, RvD2, 17(R,S)-RvD4, RvD3, RvD5, RvD5n3-DPA, RvE1, 15-epi-15-F2t-IsoP, 15-F2t-IsoP, 5-trans-PGF2α, prostaglandin (PG)F2α, 8-F2t-IsoP, 5 epi 5-F2t-IsoP/5-F2t-isoprostane (IsoP), 5(RS)-5-F2c-IsoP, 15-F3t-IsoP, 17-trans-PGF3α, thromboxane (TX)B2, PGD2, PGE2, 8-iso-PGE2, several Oxylipin MaxSpec® LC-MS Mixtures (9 eicosapentaenoic acid (EPA)-oxylipins-Mix cat#21393, 9 EPA-CYP 450 oxylipins-Mix #21394, 3 alpha-linolenic acid (ALA)/ gamma-linolenic acid (GLA) oxylipins-Mix #22638, 10 docosahexaenoic acid (DHA)-oxylipins-Mix #22280, 10 DHA-CYP 450 oxylipins-Mix #22639, linoleic acid (LA)-Mix #22638, 9 arachidonic acid (AA) oxylipins-Mix #20666) and deuterated standards: LTB4-d4, LXA4-d5, 17(R)-RvD1-d5, RvD2-d5, 5(R,S)-5-F2c-IsoP-d11, 5(R)-5-F2t-IsoP-d11, 5(S)-5-F2t-IsoP-d11, LXA4-d5, 5(S)-HETE-d8, 8-F2t-IsoP-d4, 15-F2t-IsoP-d4, 15LJepiLJ15-F2t-IsoP-d4, PGE2-d9, PGF2α-d4, TXB2-d4 were all obtained from Cayman Chemical (Ann Arbor, MI, USA). Butylated hydroxytoluene and indomethacin was acquired from Sigma-Aldrich (Oakville, ON, Canada). Acetonitrile (HPLC grade) and methanol (HPLC grade) were bought from Fisher Scientific (Ottawa, ON, Canada). Ammonium hydroxide and acetic acid were purchased from VWR International (Mont-Royal, QC, Canada) and ethanol 99% from Commercial Alcohols (Toronto, ON, Canada).

#### 2.8.2 Analysis of oxylipins

Neonatal microglia were seeded in T25 flasks (1×10^6^ cells). After 24 h treatment (control, ATGlistatin and LPS +/-ATGListatin), cells were collected with a 1:10 methanol:water solution. 30 µL of butylated hydroxytoluene (22 M) and indomethacin (625 µM) in ethanol were added to the 750 µL collected media and cells to avoid oxidation and de novo synthesis of prostaglandins. Prior to freeze-drying, 10 µl of a 25 ng/mL mixture of deuterated standards in ethanol was added to media and cells. After the drying process, cells and media were reconstituted with 0.289 mL of 41% ethanol in water. To ensure protein precipitation, 145 µL of acetonitrile was added to the reconstituted samples and then, allowed to rest for 5 minutes at -20 °C. This step was followed by centrifugation at 16060 g for 5 minutes after quick specimen thawing and mixing. Supernatants were mixed to 600 µL of 0.01 M ammonium hydroxide solution in water and loaded on conditioned solid phase extraction cartridges (SPE; Oasis MAX 60mg/3cc/30µm). SPE cartridges were washed with ammonium hydroxide solution (0.01 M) followed with acetonitrile/methanol (8:2) solution. Oxylipins elution, drying and reconstitution was achieved as described previously (Dort et al., 2021). Samples were analyzed by HPLC-MS/MS system (Bilodeau et al., 2021) with the following ternary gradient of solvents: water (A), methanol (B) and acetonitrile (C) set to 55%, 37% and 8% respectively were all prepared with 0.01% acetic acid. Solvent B was maintained for 9 min and solvent C for 2 minutes. Solvent C was further increased to 28.9% at 7.5 min and maintained to this plateau until 9 min. Then, solvent B was ramped to 90% and solvent C was decreased to 10% over 4 min, and finally kept 2 min before returning to initial proportions for 5 min to allow column re-equilibration. Statistical analysis was achieved using Metaboanalyst 5.0 in the first instance. Oxylipins that passed the threshold of significance (*p*<0.05, fold change (FC) expressed in log2 < 0.6 or > 1.5) were analysed further.

### 2.9 Untargeted lipidomics

Primary microglia (neonatal) were seeded at 2×10^6^ in 150 mm uncoated glass dishes. After 24 h treatment (control, ATGListatin, LPS +/-ATGListatin), cells were dissociated for 5 min with trypsin (0.05%) and DNAse I. Digestion was neutralized with 10% BSA in ice cold PBS, and cells were pelleted at 600 g for 10 minutes. Cells were resuspended in PBS and counted with a hemocytometer. 1.11×10^6^ cells were taken for further processing and 10% of sample taken for protein quantification. The remaining 90% were pelleted at 600 g for 10 minutes, supernatant aspirated and pellet snap-frozen in liquid nitrogen and stored at -80 °C until analysis.

Lipid extraction, sample and data analysis were performed using a previously validated untargeted lipidomic workflow (Forest et al., 2018; Lefort et al., 2023; Ruiz et al., 2019). In brief, lipids were extracted from microglia pellets and spiked with six internal standards: LPC 13:0, PC19:0/19:0, PC14:0/14:0, PS12:0/12:0, PG15:0/15:0, and PE17:0/17:0 (Avanti Polar Lipids). Protein concentration was determined using BioRad Bradford dye-binding method as described in section 2.4. Samples were injected (from 1.66 to 9.92uL according to protein concentration) into a 1290 Infinity high resolution HPLC coupled with a 6530 Accurate Mass quadrupole time-of-flight (LC-QTOF) (Agilent Technologies) via a dual electrospray ionization (ESI) source. Elution of lipids was assessed on a Zorbax Eclipse plus column (Agilent Technologies) maintained at 40LJ°C using a 83 min chromatographic gradient of solvent A (0.2% formic acid and 10LJmM ammonium formate in water) and B (0.2% formic acid and 5LJmM ammonium formate in methanol/acetonitrile/methyl tert-butyl ether [MTBE], 55:35:10 [v/v/v]). Data acquisition was performed in positive ionisation mode. All samples were processed and analyzed as a single batch. MS quality controls (QCs) were performed by (i) injecting 3 “in-house” QC samples and blanks at the beginning, middle and end of the run and (ii) monitoring the six internal standards spiked in samples for signal intensity, mass mass-to-charge ratios (m/z) and retention time (RT) accuracies. Mass spectrometry (MS) raw data processing was achieved as previously described using Mass Hunter B.06.00 (Agilent Technologies) for peak picking and an in-house bioinformatic script (Forest et al., 2018) for data processing. The resulting final dataset included 1296 high-quality MS signals, or features, defined by their m/z, RT and signal intensity and for a same unique lipid, several adducts referred as features may be annotated. Lipid annotation was achieved by MS/MS analysis for the most significant lipid features that passed the following threshold of significance (using a 2-tailed unpaired Student’s t test): *p* < 0.05 and a fold change (FC) expressed in log2 < 0.8 or > 1.25 for the comparisons (i) LPS vs. DMSO, (ii) ATGLi vs. DMSO, and (iii) ATGLi + LPS vs. LPS. In addition, we used data alignment with our in-house database, which contains > 500 unique lipids with previously determined MS/MS spectra, for the remaining features. Side chains are indicated when annotated using MS/MS with certitude. When not identified, we only mentioned the total number of carbons and double bounds.

### 2.10 Statistics

Data is presented as mean ± S.E.M. Data prepared for analysis in Excel and analysed and prepared for presentation using GraphPad Prism (v9.0.0). Intergroup comparisons were performed using Two-way ANOVAs with post-hoc Tukey’s tests, One-way ANOVAs with post-hoc Tukey’s or Student’s t-test’s as appropriate. *P* < 0.05 was considered significant.

## 3 Results

### 3.1 Pan-lipase inhibition by ORlistat reduced LPS-induced inflammation in primary microglia

To establish whether LDs accumulate in LPS-activated primary microglia, as previously reported by others (Khatchadourian et al., 2012; Li et al., 2023; Loving et al., 2021; Marschallinger et al., 2020), primary mouse microglia were treated with LPS (0.1 µg/mL) for 6 h. LPS increased accumulation of LDs from a mean of 2.1 to 9.9 LDs per cell (363%, *p* = 0.0012; Fig. **1A-B**). To determine whether LPS-induced LD accumulation could result from a decrease of TG lipolysis, the expression of the main glycerolipid lipases and proteins that regulate lipase activity were measured by qPCR. ATGL mRNA level was downregulated by LPS after 6 h (39%, *p*<0.0001; Fig. **1C**). This downregulation of ATGL mRNA translated into reduced protein level (68%, *p* = 0.0268, Fig. **1D-E**). Expression of Hormone Sensitive lipase (HSL), a TG and DAG lipase, was not affected by LPS (*p* = 0.2021, Fig. **1C**). Expression of co-regulators of ATGL activity, PLIN2 and ABHD5 (CGI-58) was not regulated at the transcriptional level by LPS (*p* = 0.32 & *p* = 0.053 respectively, Fig. **1C**), whereas PLIN3 was downregulated by 53% (*p*<0.0001, Fig. **1C**).

**Fig 1.**
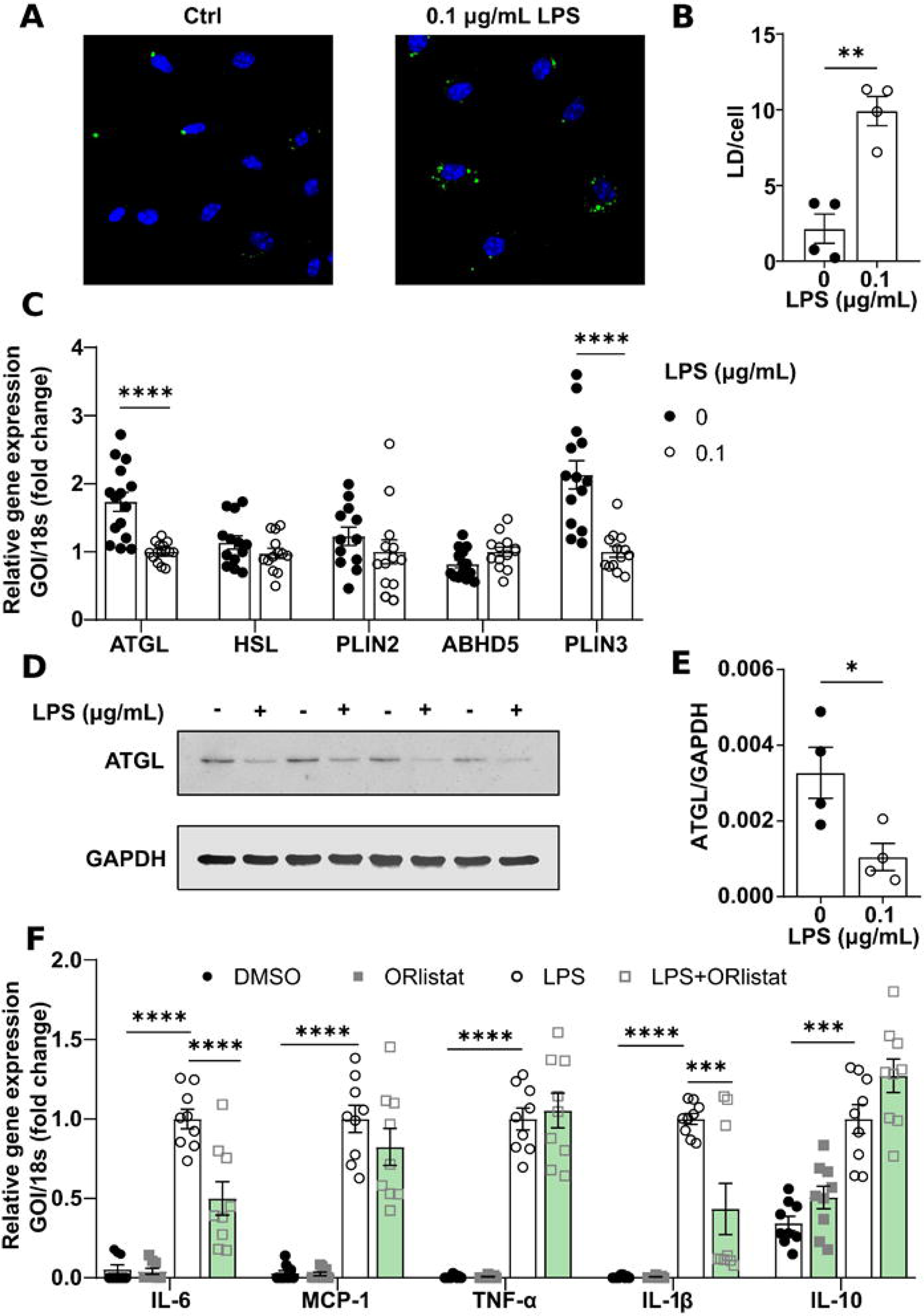
Lipase inhibition by ORlistat reduces LPS-induced neuroinflammation in primary microglia. **A.** Representative images of primary microglia treated 6 h with 0.1 µg/mL LPS and stained with BODIPY and Hoescht. **B.** Average number of lipid droplets (LD) per cell (n=4, 5 cells/n). **C.** Relative gene expression of lipases and co-regulators (n=12-15). **D.** Representative immunoblot and **E.** densitometric analysis of expression of ATGL protein (n=4). * *p*<0.05; ** *p*<0.01; **** *p*<0.0001, Student’s t-test. **F.** Relative gene expression of cytokines after 6 h LPS ± 50 µM Orlistat (n=9). *** *p*<0.001; **** *p*<0.0001; Two-way ANOVA with post-hoc Tukey.

To investigate whether TG lipolysis regulates pro-inflammatory responses in microglia, the expression of cytokines was measured by qPCR in primary microglia derived from pups treated for 6 h with LPS with or without the pan-lipase inhibitor ORlistat (50 µM). ORlistat did not affect the basal expression of cytokines but attenuated LPS-induced IL-6 gene expression by 50% (Tukey’s test, *p*<0.0001, q = 7.952, dF = 32; Fig. **1F**) and IL-1β by 57% (Tukey’s test, *p* = 0.0002, q = 6.880, dF = 32; Fig. **1F**). MCP-1 (Tukey’s test, *p* = 0.33, q = 2.416, dF = 32; Fig. **1F**) and TNF-α (Tukey’s test, *p* = 0.94, q = 0.8, dF = 32; Fig. **1F**) expression were not affected by ORlistat compared to the LPS. Expression of the anti-inflammatory cytokine IL-10 induced by LPS was not significantly affected by ORlistat (Tukey’s test, *p* = 0.10, q = 3.339, dF = 32; Fig. **1F**). These findings demonstrate that TG lipolysis is required for microglial upregulation of specific cytokines in response to LPS.

### 3.2 Pharmacological inhibition of ATGL activity blunted LPS-induced inflammatory responses in primary microglia

Since ATGL but not HSL expression was downregulated by LPS, we hypothesised that ATGL may be the main regulator of microglial LDs and effector of reduced inflammation in response to pan-lipase inhibition by Orlistat. In line with results in Fig. **1A**, LPS induced LD accumulation (267% *p* = 0.0442, q = 3.850, dF = 44, Fig. **2A**) in a distinct set of experiments on primary microglia. Inhibition of ATGL with the specific inhibitor ATGListatin led to a similar increase in LD number (280%, *p* = 0.0318, q = 4.042, dF = 44, Fig. **2A**). No additive effect of LPS and ATGListatin were observed on LD number (33%, *p* = 0.6080, q = 1.747, dF = 44, Fig. **2A**). To investigate whether ATGL inhibition is sufficient to recapitulate the effect of pan-lipase inhibition (Fig. **1F**), primary neonatal microglial cultures were treated for 6 h with ATGListatin (50 µM) and/or LPS. In neonatal cultures, inhibition of ATGL decreased LPS-induced IL-6 (46%, Student’s t-test, *p*<0.0001; Fig. **2B**), MCP-1 (35%, Student’s t-test, *p* = 0.02; Fig. **2B**) and IL-1β (38%, Student’s t-test, *p*<0.0001; Fig. **2B**) expression. In primary adult microglial cultures, ATGL inhibition recapitulated the majority of the effects observed in neonatal cultures, with ATGListatin treatment decreasing IL-6, MCP-1 and IL-1β by 44% (*p* = 0.003), 44% (*p* = 0.0189) and 38% (*p*<0.0026) respectively in LPS-treated conditions (Fig. **2C**). Importantly, these changes led to a decrease in LPS-induced cytokine secretion in neonatal microglia, most strongly observed 12-24 h post-treatment (Fig. **2D-E**). IL-6 secretion was decreased by 69% (12 h: *p* = 0.011, q = 2.13, dF = 152; 24 h: *p*<0.0001, q = 5.878, dF = 152; Fig. **2D**), while MCP-1 secretion was decreased by 62%-48% at 12 h (*p*<0.0001, q = 8.044, dF = 204) and 24 h (*p*<0.0001, q = 7.935, dF = 204); Fig. **2E**) in ATGListatin + LPS conditions vs. LPS conditions. ATGL inhibition also enhanced the LPS-induced expression of the anti-inflammatory cytokine IL-10 by 55% (*p* = 0.0097; Fig. **2B**). However, this effect was not recapitulated in adult microglia cultures, in which ATGListatin induced a 50% decrease in IL-10 expression (*p* = 0.0241; Fig. **2C**). These data suggest that inhibition of ATGL activity recapitulated in most part the effect of ORlistat and was sufficient to reduce LPS-induced pro-inflammatory responses. However, the effect of ATGL inhibition did not affect LPS-induced expression of TNF-α (*p* = 0.3 Fig. **2B-C**) suggesting that ATGL specifically regulates a subset of cytokines.

**Fig 2.**
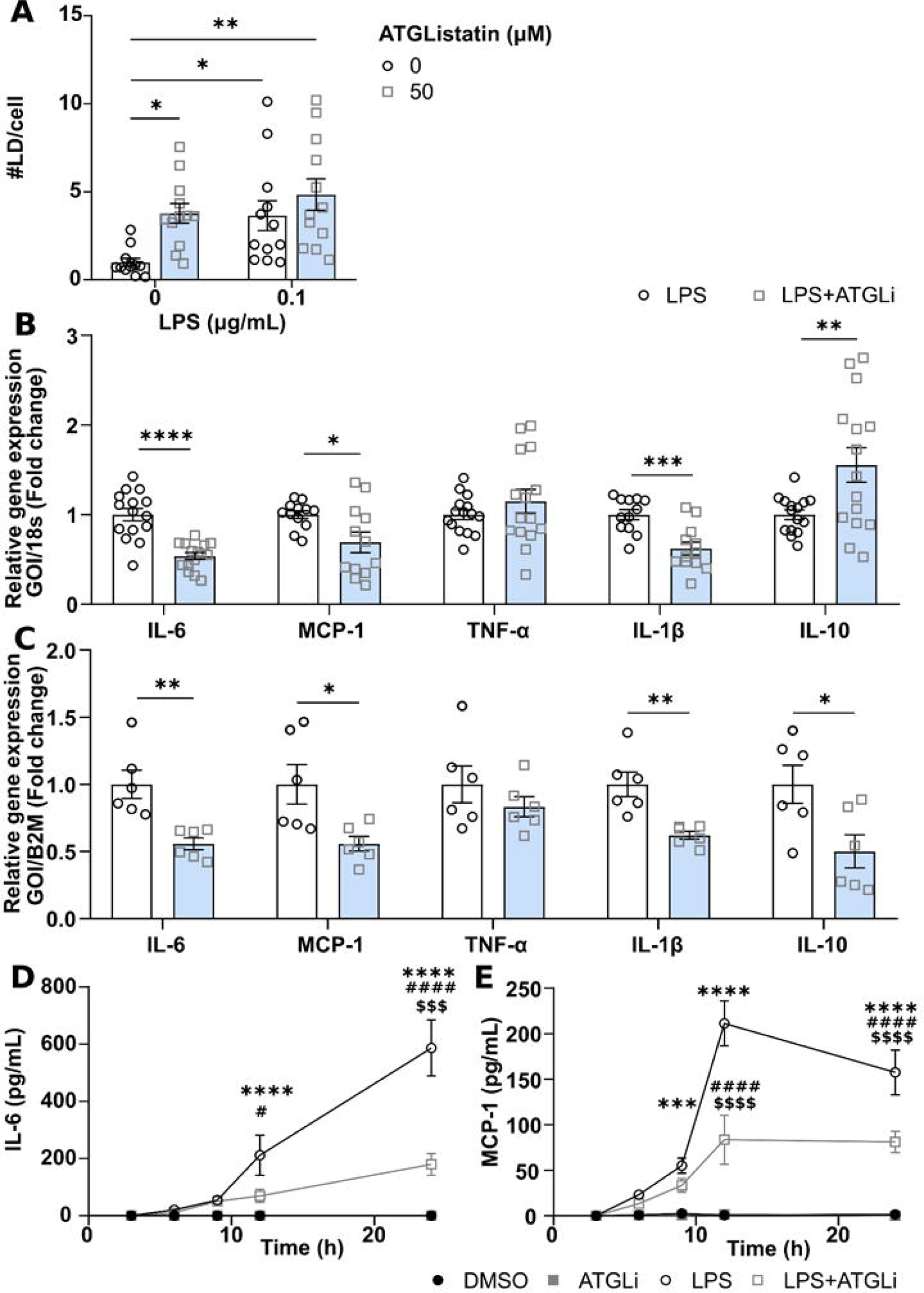
ATGL inhibition reduces LPS-induced neuroinflammation in primary microglia. Effect of treatment with 0.1 µg/mL LPS ± 50 µM ATGListatin on LD and cytokines. **A.** Average number of lipid droplets (LD) per cell (n=4, 3 fields of view/n). * *p*<0.05; ** *p*<0.01; One-way ANOVA with post-hoc Tukey. Relative gene expression of cytokines in **B.** neonatal microglia (n=15) and **C.** adult male microglia (n=6) in response to a 6 h LPS treatment. * *p*<0.05; ** *p*<0.01 *** *p*<0.001; **** *p*<0.0001; Two-way ANOVA with post-hoc Tukey. Extracellular cytokine concentration of **D.** IL-6 **E.** MCP-1 from neonatal microglia up to 24 h after treatment. n=10-15; (* LPS vs ctrl; # LPS vs ATGListatin + LPS; $ ATGLi vs ATGLi + LPS); * *p*<0.05; *** *p*<0.001; **** *p*<0.0001; Two-way ANOVA with post-hoc Tukey.

ATGL activity releases fatty acids which can be used as substrates for fatty acid oxidation (FAO) in the mitochondria. To assess whether the action of ATGListatin on LPS responses could be dependent on reduced FAO, we assessed whether ATGListatin decreases FAO and whether FAO inhibition with etomoxir (10 µM) recapitulates the effect of ATGL inhibition in neonatal microglia. ATGListatin and etomoxir reduced FAO by 33% (*p* = 0.0228 q = 4.109, dF = 20, Supplementary Fig. 1) and 49% (*p*<0.0001, q = 7.958, dF = 20, Supplementary Fig. 1), respectively. However, etomoxir did not affect the expression and secretion of MCP-1 or IL-6 in response to LPS (Supplementary Fig. 1).

### 3.3 Inhibition of ATGL reduced the release of prostanoids induced by LPS

Oxylipins are lipid mediators of inflammation derived from polyunsaturated fatty acids that can be released by LDs lipolysis. We investigated the impact of ATGL inhibition on intracellular and extracellular levels of oxylipins in neonatal microglia treated for 24 h with ATGListatin and/or LPS. Among 88 oxylipin species analysed by targeted LC-MS/MS, 79 had detectable levels. The overall changes in detected oxylipins are shown as heatmaps in Supplementary Fig. 2. Using principle component analysis (PCA) no samples were identified as outliers. A partial separation between the LPS condition and all other conditions was observed in the culture media (Fig. **3A**). Partial separation was also observed between the vehicle and ATGListatin conditions, suggesting that inhibition of ATGListatin altered the release of oxylipin species at baseline. 95% of the variance between groups was explained by component 1, and 4.1% of the remaining variance was explained by component 2. For intracellular oxylipins, no separation between groups was observed using PCA (Fig. **3D**). Volcano plot analysis was performed to identify lipid species significantly affected by LPS vs. controls (Fig. **3B, E**), and by ATGListatin + LPS compared to LPS (Fig. **3C, F**). FC > 1.5 and < 0.6 and *p <* 0.05 was used as the threshold to identify significant changes in oxylipin levels. Three prostanoids were significantly upregulated by LPS and reduced by co-treatment with ATGListatin in the media: prostaglandin E2 (PGE_2_), D2 (PGD_2_), and thromboxane B2 (TXB_2_). PGF_2a_ level in the intracellular fraction was regulated in a similar manner (ctrl vs LPS, Fig. **3E**; LPS vs ATGListatin + LPS, Fig. **3F**). Two-way ANOVA analysis revealed that inhibition of ATGL decreased LPS-induced PGE_2_ release by 81% (Tukey’s test, *p*<0.0001, q = 8.482, dF = 20, Fig. **4A**), PGD_2_ by 82% (Tukey’s test, *p* = 0.0002, q = 7.441, dF = 20, Fig. **4A**), and TXB_2_ by 71% (Tukey’s test, *p* = 0.0003, q = 7.118, dF = 20, Fig. **4A**). The intracellular level of PGF_2a_ was reduced by 51% (Tukey’s test, *p* = 0.0023, q = 5.940, dF = 20, Fig. **4B**).

**Fig 3.**
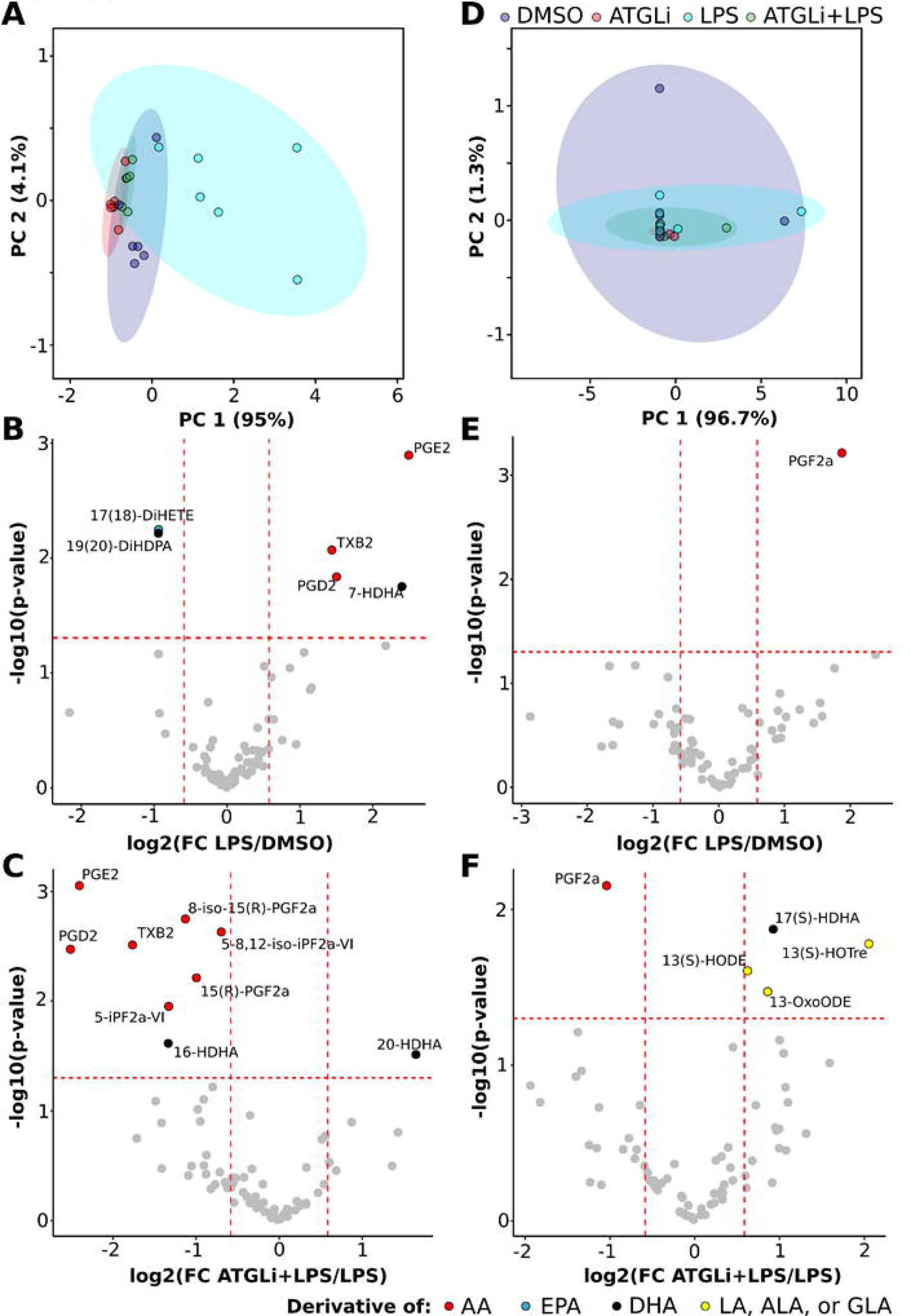
Inhibition of ATGL alters the release of oxylipins. Oxylipins analysis by LC-MS/MS in response to 24 h treatment with 0.1 mg/mL LPS ± 50 µM ATGListatin. PCA analysis of **A.** extracellular and **D.** intracellular oxylipins. Volcano plot comparing LPS vs. vehicle in the **B.** extracellular and **E.** intracellular fraction. Volcano plot comparing ATGListatin + LPS vs. LPS in the **C.** extracellular and **F**. intracellular fraction. FC=1.5, *p*<0.05.

**Fig 4.**
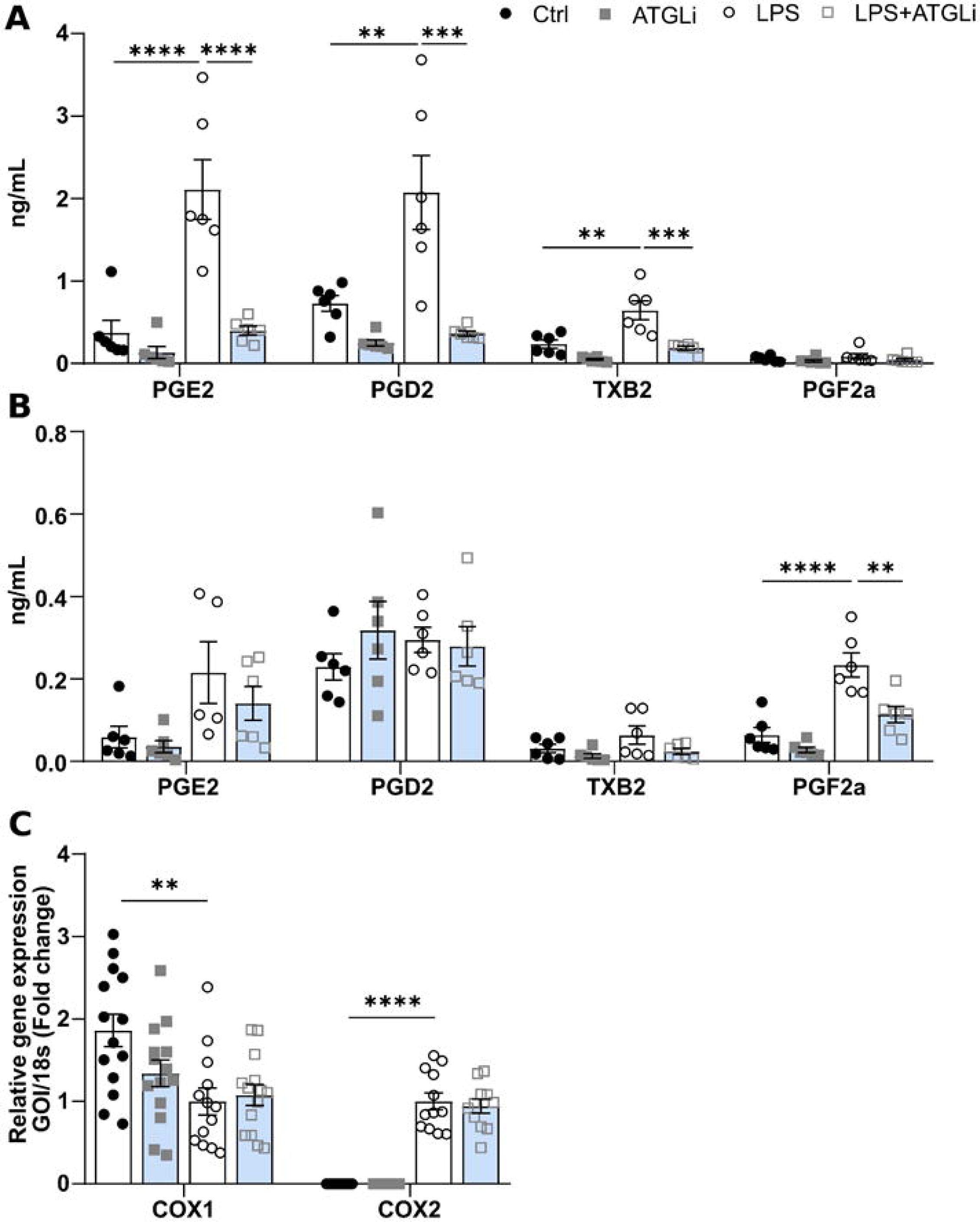
Inhibition of ATGL alters prostanoid release independently from COX1/2 expression. Effect of 24 h 0.1 mg/mL LPS ± 50 µM ATGListatin on oxylipins identified by LC-MS/MS. Altered oxylipins in the **A.** extracellular and **B.** intracellular fractions (n=6). **C.** Gene expression of COX-1 and COX-2 6 h post LPS ± ATGListatin treatment (n=15). * *p*<0.01; *** *p*<0.001; **** *p*<0.0001; Two-way ANOVA with post-hoc Tukey.

The oxylipins affected by ATGListatin are generated from arachidonic acid via the enzymes cyclooxygenase 1/2 (COX-1/2). To assess whether decreased oxylipin levels were related to changes in enzyme expression COX-1/2 expression was measured by qPCR in neonatal microglia treated with ATGListatin and/or LPS. While LPS induced the expected downregulation of COX-1 (Tukey’s test, *p* = 0.003, q = 5.208, dF = 51; Fig. **4C**), and upregulation of COX-2 (*p*<0.0001, q = 15.36, dF = 43; Fig. **4C**), ATGL inhibition did not alter these responses (COX-1 downregulation, *p* = 0.0067, q = 4.832, dF = 51, Fig. **4C**; COX-2 upregulation, *p*<0.0001, q = 14.15, dF = 43; Fig. **4C**).

### 3.4 ATGL inhibition altered ceramide species in neonatal microglia

Lipid-derived molecules are important to many aspects of the inflammatory response beyond production of cytokines and prostanoids. To assess more broadly the impact of ATGL activity on the microglial lipidome, untargeted lipidomic profiling was performed by LC-MS/MS on primary cells treated for 24 h with LPS and/or ATGListatin. Considering *p* < 0.05 and FC > 1.25 and < 0.8 as significant for the identified lipid species, we found 4 lipids significantly affected by LPS vs. DMSO conditions (Fig. **5A**), and 5 lipids regulated by ATGListatin vs. DMSO (Fig. **5B**). Four lipids were modulated by ATGListatin + LPS vs. LPS conditions (Fig. **5C**). These lipids were predominantly ceramides. Three ceramide species were significantly upregulated by LPS Cer(d18:0/16:0), *p* = 0.0186, FC = 2; Cer(d40:0), *p* = 0.0454, FC = 1.81; Cer(d18:0/24:0), *p* = 0.0111, FC = 1.72; Table 2). Co-treatment with ATGListatin appeared to exacerbate the effect of LPS, further upregulating most of the identified ceramide species in the same direction as LPS treatment alone. A similar pattern was observed for sphingomyelin (d18:0/18:0) which was increased by ATGListatin + LPS. However, Cer(d18:1/24:0) (*p* = 0.0156, FC = 1.8) and Cer(d40:1) (*p* = 0.0289, FC = 1.8) were significantly downregulated by co-treatment with ATGListatin + LPS vs. LPS, whereas LPS non-significantly upregulated these species (Table 2). The overall changes (significant and non-significant) in identified ceramides, sphingomyelins and triglycerides are shown as heatmaps in Supplementary Fig. 3.

**Fig 5.**
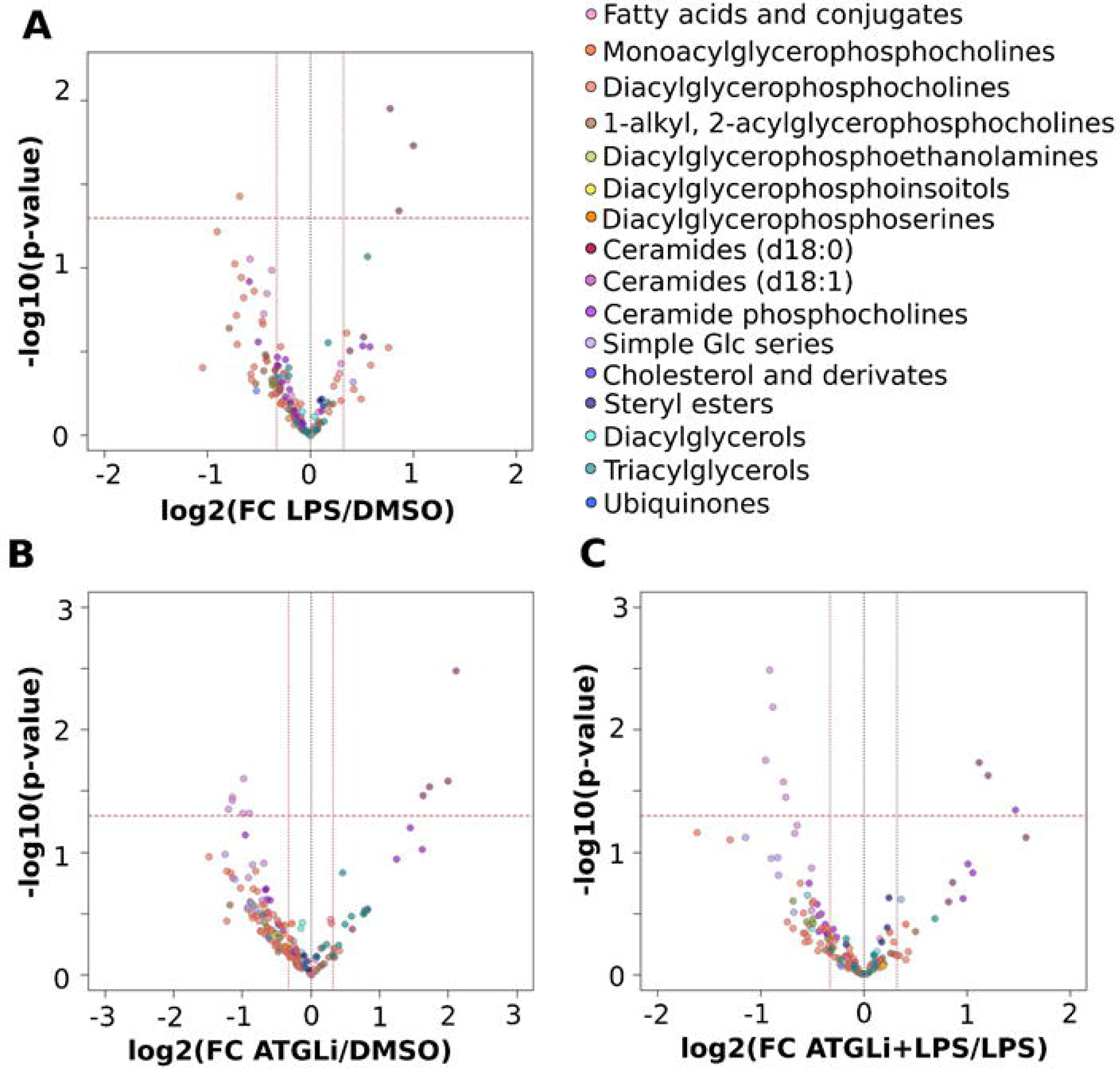
Inhibition of ATGL alters microglial ceramide profile. Effect of 24 h 0.1 mg/mL LPS ± 50 µM ATGListatin on the lipidome as identified by LC-MS/MS. Volcano plots showing 162 MS signals or lipid features (154 unique lipids), represented by a dot and defined by RT, m/z and signal intensity, annotated either by alignment with our in-house database to specific lipid entities (145 features; 142 unique lipids) or by MS/MS (17 features; 12 unique lipids). Coloured dots referring to the various lipid subclasses are indicated in the legend. Features were defined as significantly different with a *p* < 0.05, FC > 1.25. **A.** LPS vs. DMSO **B.** ATGListatin vs. DMSO and **C.** ATGListatin and LPS vs. LPS.

**Table 2.** Ceramides and sphingomyelin altered by LPS and/or ATGListatin. Effect of LPS (0.1 µg/mL) ± ATGListatin (50 µM) on microglial ceramides (Cer) and shingomyelin (SM) identified by untargeted LC-MS/MS. Data are represented as absolute (abs) fold change (FC), with column indicating upregulation or downregulation. *P* < 0.05, FC > 1.25 considered as significant upregulation (red) or downregulation (blue).

|  | LPS vs Ctrl |  |  | ATGLi vs Ctrl |  |  | ATGLi+LPS vs LPS |  |  |
| --- | --- | --- | --- | --- | --- | --- | --- | --- | --- |
| Annotated ID | p | FC (abs) |  | p | FC (abs) |  | p | FC (abs) |  |
| Cer(d18:0/16:0) [M+H]<sup>+</sup> | **0.0186** | 2.00 | up | 0.4183 | 1.52 | up | 0.1317 | 1.82 | up |
| Cer(d18:0/24:0) [M+H]<sup>+</sup> | **0.0111** | 1.72 | up | **0.0033** | 4.33 | up | **0.0124** | 2.17 | up |
| Cer(d18:0/24:0) [M+H+44]<sup>+</sup> | 0.3103 | 1.30 | up | **0.0344** | 3.11 | up | **0.0142** | 2.31 | up |
| Cer(d18:0/24:1) [M+H]<sup>+</sup> | 0.2577 | 1.43 | up | **0.0262** | 3.99 | up | **0.0422** | 2.98 | up |
| Cer(d40:0) [M+H]<sup>+</sup> | **0.0454** | 1.81 | up | **0.0290** | 3.31 | up | 0.1987 | 1.77 | up |
| Cer(d18:1/24:0) [M+H]<sup>+</sup> | 0.1033 | 1.30 | down | **0.0252** | 1.97 | down | **0.0033** | 1.88 | down |
| Cer(d18:1/24:0) [M+H-H<sub>2</sub>O]<sup>+</sup> | 0.1422 | 1.34 | down | **0.0352** | 2.21 | down | **0.0094** | 1.84 | down |
| Cer(d18:1/24:0) [M+H+44]<sup>+</sup> | 0.1876 | 1.37 | down | **0.0376** | 2.21 | down | **0.0369** | 1.69 | down |
| Cer(d18:1/24:1) [M+H]<sup>+</sup> | 0.7734 | 1.06 | down | **0.0482** | 1.99 | down | 0.0700 | 1.59 | down |
| Cer(d18:1/24:1) [M+H-H<sub>2</sub>O]<sup>+</sup> | 0.5726 | 1.14 | down | **0.0445** | 2.30 | down | 0.0659 | 1.56 | down |
| Cer(d18:1/24:1) [M+H]<sup>+</sup> | 0.7734 | 1.06 | down | **0.0482** | 1.99 | down | 0.0700 | 1.59 | down |
| Cer(d40:1) [M+H-H<sub>2</sub>O]<sup>+</sup> | 0.8959 | 1.03 | up | 0.1264 | 1.80 | down | **0.0289** | 1.72 | down |
| Cer(d40:1) [M+H+44]<sup>+</sup> | 0.3737 | 1.23 | up | 0.1221 | 1.61 | down | **0.0156** | 1.93 | down |
| Cer(d41:1) [M+H]<sup>+</sup> | 0.4512 | 1.27 | down | 0.2453 | 1.63 | down | 0.2988 | 1.42 | down |
| Cer(d42:2) [M+H-H<sub>2</sub>O]<sup>+</sup> | 0.0879 | 1.50 | down | 0.1605 | 1.87 | down | 0.3206 | 1.34 | down |
| SM(d18:0/18:0) [M+H]<sup>+</sup> | 0.2939 | 1.50 | up | 0.0946 | 3.08 | up | **0.0448** | 2.78 | up |

### 3.5 Microglia specific ATGL loss-of-function reduces LPS-induced inflammation and sickness behaviour in male mice

To determine the role of ATGL on the microglial inflammatory response *in vivo* we generated a novel mouse model allowing a specific and inducible knock out of ATGL in microglia in adult mice using the Cx3CR1-CreER-YFP strain (CX3CR1^ATGL^ WT and KO, see methods). To validate loss of ATGL specifically from microglia, fluorescence activated cell sorting was performed 4 weeks after tamoxifen administration in adult male mice. Relative expression of IBA1 and ATGL was measured by qPCR in YFP^+^/CD11b^+^ (microglial/macrophage) and YFP^-^/CD11b^-^ (non-microglial) cell populations from brain and spleen samples (Supplementary Fig. 4). As expected, IBA1 was expressed in the YFP^+^/CD11b^+^ population and undetectable in the YFP^-^ cells. IBA1 expression was not affected in CX3CR1^ATGL^ KO YFP^+^/CD11b^+^ brain cells (*p*=0.31). In YFP^+^ cells, ATGL expression was decreased by 95% (*p*=0.017) in CX3CR1^ATGL^ KO mice compared to control Cx3CR1-CreER-YFP littermates (CX3CR1^ATGL^ WT). ATGL expression was not affected in YFP^-^ brain cells (Supplementary Fig. 4B) and CD11b+ spleen cells (Supplementary Fig. 4D) thereby confirming that loss of ATGL expression was restricted to microglia. Loss of microglial ATGL did not affect the gross phenotype of male mice, including body weight or anxiety-like behaviour in basal conditions measured by elevated plus maze (EPM) and open field (OF) tests (Supplementary Fig. **4 E-J**).

To assess whether loss of ATGL in microglia affects pro-inflammatory responses induced by LPS, male mice were administered *i.p.* with LPS (0.83 mg/kg) and cytokine expression was measured 3 h after in different brain regions as we previously described (Demers et al., 2020). As expected, LPS increased the expression of IL-6 and MCP-1 in all regions analysed including the cortex, hippocampus and mediobasal hypothalamus (MBH) of male CX3CR1^ATGL^ WT mice (Fig. **5**). Consistent with our observation *in vitro*, loss of microglial ATGL expression decreased LPS-induced IL-6 expression in the MBH (57%, Tukey’s test, *p* = 0.0081, q = 5.084, dF = 22; Fig. **6A**). Although not statistically significant, there was a trend towards reduced LPS-induced IL-6 and MCP-1 in the other brain regions of CX3CR1^ATGL^ KO male mice (Fig. **6B-C**). A similar pattern was also observed in mice injected with 2 lower doses of LPS (0.5 mg/kg). In these animals, LPS-induced IL-6 expression was significantly reduced in the cortex (52%, *p* = 0.0005; Fig. **6E**) and the MBH (53%, *p* = 0.0035; Fig. **6D**) but not in the hippocampus (*p* = 0.25; Fig. **6F**). LPS-induced expression of MCP-1 across brain regions was not affected in CX3CR1^ATGL^ KO as compared to CX3CR1^ATGL^ WT mice (Fig. **6**).

**Fig 6.**
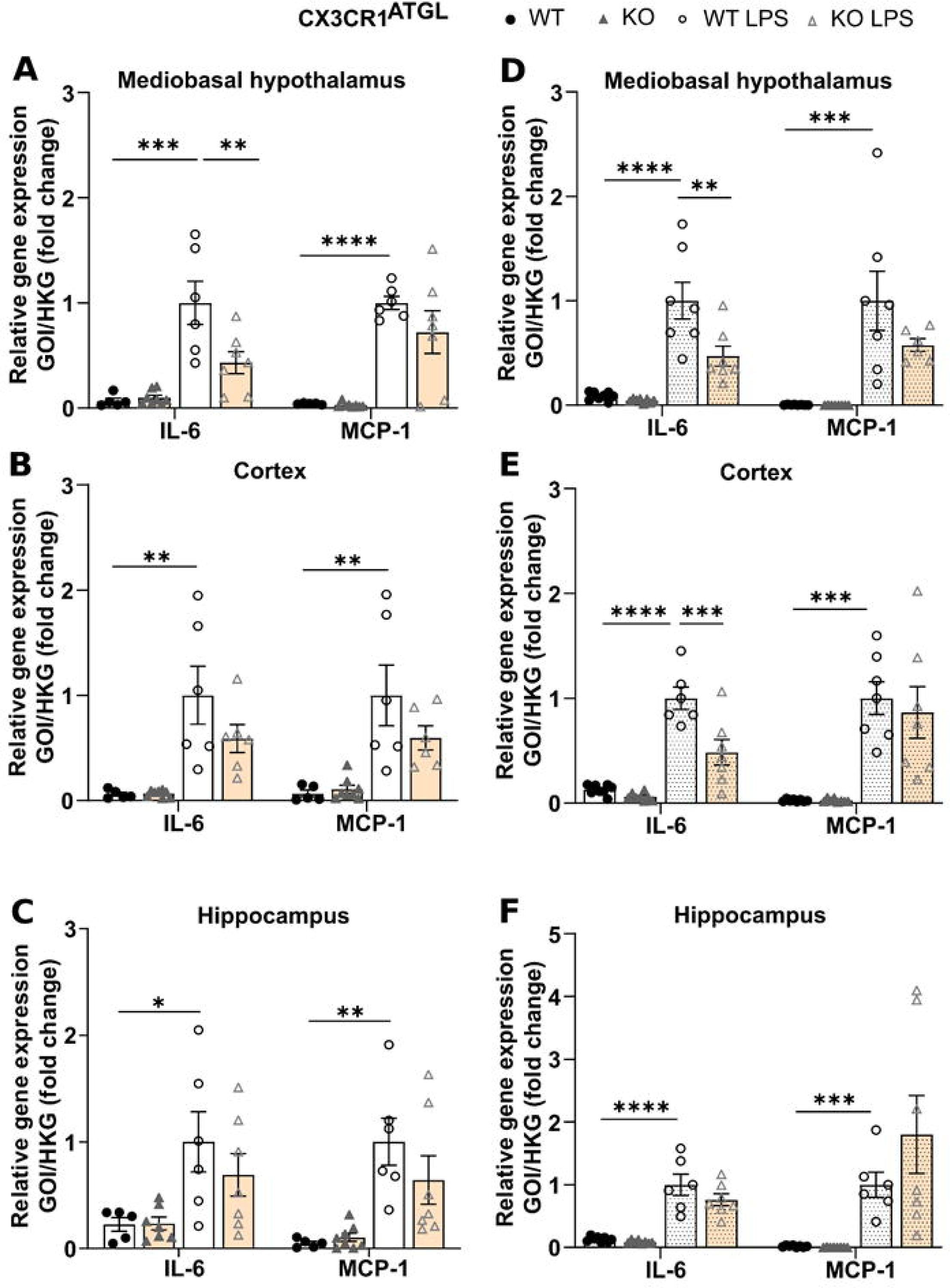
ATGL deletion in microglia blunts LPS-induced neuroinflammation in a brain region-dependent manner. Impact of microglial ATGL loss-of-function (CX3CR1^ATGL^KO) on LPS-induced cytokine expression in mouse brain regions compared to Cre expressing male littermates (CX3CR1^ATGL^WT). Relative gene expression in the **A.** mediobasal hypothalamus, **B.** cortex and **C.** hippocampus, 3 h after 0.83 mg/kg LPS or saline *i.p.* (n=6-8). Relative gene expression in the **D.** mediobasal hypothalamus, **E.** cortex and **F.** hippocampus, 3 h after 2 × 0.5 mg/kg LPS or saline *i.p.* 24 h apart (n=7-8). * *p*<0.05; ** *p*<0.01; *** *p*<0.001; **** *p*<0.0001; Two-way ANOVA with post-hoc Tukey.

Neuroinflammation is well known to contribute to psychomotor and emotional disturbances induced by systemic inflammation of infection. To establish whether microglial ATGL affects sickness-and anxiety-like behaviours induced by LPS, mice were injected *i.p.* with saline or LPS (single dose, 0.83 mg/kg) and were subjected to EPM and light-dark box (LDB) tests 12 h after the injection. In agreement with EPM results presented in Supplementary Fig. 4E-G in naïve mice (no injection), distance, entries and time in open arms were similar in CX3CR1^ATGL^ WT and KO male mice treated with saline. In CX3CR1^ATGL^ WT mice, LPS reduced locomotor activity by 28% during the EPM compared to control littermates injected with saline (Tukey’s test, *p* = 0.0099, q = 4.755, dF = 34, Fig. **7A**). LPS-induced hypo-locomotion was abrogated in CX3CR1^ATGL^ KO animals (5%, Tukey’s test, *p* = 0.07, q = 1.05, dF = 34, Fig. **7A**). Time and number of entries in the open arms were not affected by LPS in agreement with our previous study (Demers et al., 2020). During the LDB test, distance was reduced in saline-treated CX3CR1^ATGL^ KO male mice vs. control littermates. While LPS led to hypo-locomotion in control littermates (48%, Tukey’s test, *p* = 0.0032, q = 5.462, dF = 28, Fig. **7D**) as well as reduced time (47%, Tukey’s test, *p* = 0.0169, q = 4.928, dF = 28, Fig **7E**) and number of entries (54%, Tukey’s test, *p* = 0.0068, q = 5.051, dF = 28, Fig. **7F**) in the light zone compared to saline controls, these responses were absent in CX3CR1^ATGL^ KO male mice (Tukey’s test; *p* = 0.9904, q = 0.4247, dF = 28, Fig. **7D**; *p*=0.45 q = 2.134, dF = 28, Fig. **7E**; *p* = 0.2117, q = 2.830, dF = 28, Fig. **7F**). When the data from both tests were combined into a measure of emotionality (z-score) as described by Guilloux *et al*. (2011), we found that LPS strongly reduced the z-score (Tukey’s test; *p* = 0.0002, q = 6.876, dF = 27, Fig. **7G**) in control littermates while it was not affected in CX3CR1^ATGL^ KO mice (Tukey’s test; *p* = 0.3216, q = 2.466, dF = 27, Fig. **7G**).

**Fig 7.**
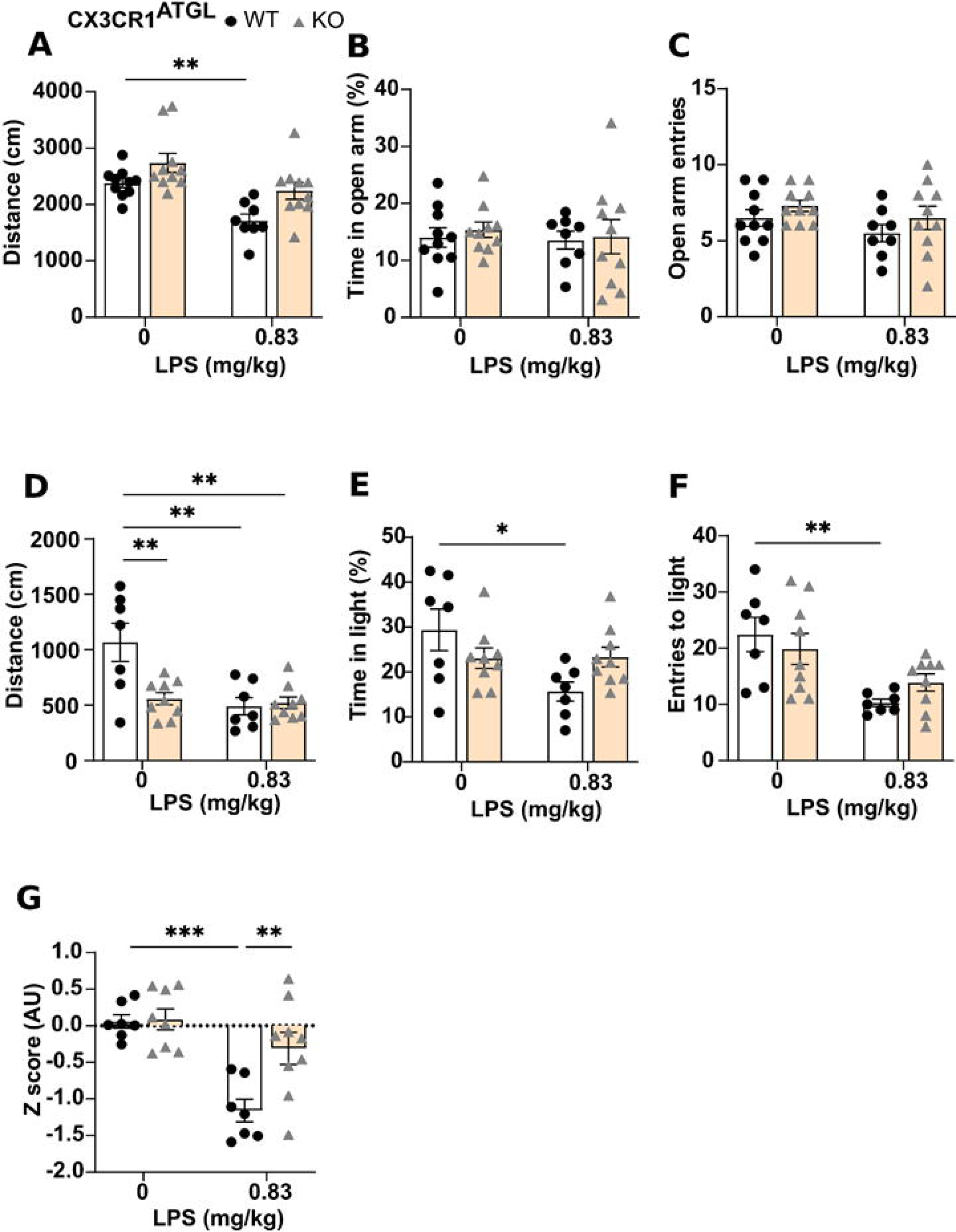
Loss of ATGL from microglia alters the behavioural response to LPS in male mice. Impact of microglial ATGL loss-of-function (CX3CR1^ATGL^KO) on LPS-induced sickness- and anxiety-like behaviours vs. CX3CR1^ATGL^WT male littermates (12 h post 0.83 mg/kg LPS or saline *i.p.*) (n=8-10). Measures of **A.** distance travelled, **B.** percentage of total time spent in open arms, and **C.** open arm entries in elevated plus maze in 5 minutes. Measures of **D.** distance travelled **E.** percentage of total time spent in light, and **F.** entries in the light section of a light-dark box in 5 minutes. **G.** Emotionality score. * *p*<0.05; ** *p*<0.01; *** *p*<0.001; **** *p*<0.0001; Two-way ANOVA with post-hoc Tukey.

## Discussion

Despite accumulating evidence showing that TG metabolism and LD dynamics in immune cells contribute to cell function and overall regulation of inflammation, the role of ATGL and LD lipolysis in microglia activation, neuroimmune and behavioral responses to inflammatory insult remains elusive. The present study demonstrates that ATGL regulates basal LD lipolysis in primary microglia and that ATGL inhibition in both neonatal and adult primary microglia blunts the expression and secretion of pro-inflammatory cytokines in response to LPS. Importantly, genetic deletion of ATGL specifically in microglial cells of the mouse brain alleviated pro-inflammatory cytokine responses to LPS and was sufficient to attenuate anxiety- and sickness-like behaviours induced by systemic LPS administration. At the cellular level, inhibition of FAO in primary microglia did not recapitulate the anti-inflammatory effect of ATGL inhibition while lipidomic profiling showed that ATGListatin robustly reduced the release of pro-inflammatory prostanoids and altered the balance between dihydroceramides and ceramides. Collectively, our results suggest that ATGL-dependent LD lipolysis in microglia mitigates acute neuroimmune and behavioural responses to LPS by reducing the generation of fatty acid-derived signals of inflammation.

Consistent with previous findings in primary microglia (Li et al., 2023; Marschallinger et al., 2020) or microglial cell lines (Khatchadourian et al., 2012; Loving et al., 2021), LPS treatment led to LD accumulation. Although the pathway(s) promoting microglial LD expansion remains unclear, LPS induced a robust decrease of ATGL mRNA and protein levels, without affecting HSL expression. This suggests LPS-induced LD accumulation result from decreased ATGL-dependent lipolysis. In line with this, we found that ATGL inhibition was sufficient to increase LD number in unstimulated conditions suggesting a constant lipolytic activity in microglia. Khatchadourian *et al*. (Khatchadourian et al., 2012) attributed LPS-induced LD accumulation to increased expression of the regulatory protein PLIN2 which excludes ATGL from binding LD membrane. Although PLIN2 mRNA level was not affected by LPS in our study, PLIN3 expression was strongly reduced by LPS which may contribute to decreased lipolysis (Itabe et al., 2017). The observed LPS-induced downregulation of ATGL and anti-lipolytic action promoting LD accumulation in microglia are consistent with previous studies in macrophages (Feingold et al., 2012; Huang et al., 2014). Altogether, these data suggest that LPS promotes microglial LD accumulation by reducing ATGL-dependent lipolysis. Additional work will be needed to identify the signalling and molecular pathway(s) by which LPS downregulates ATGL expression.

Our studies using the pan-triglyceride lipase (ORlistat) or the specific inhibitor of ATGL (ATGListatin) consistently demonstrated that TG lipase inhibition blunted inflammatory responses in primary microglia. ORlistat is mostly known as a weight loss therapy that prevents intestinal fat absorption by inhibiting gastric and pancreatic lipase activity. However, it was recently reported that ORlistat efficiently inhibits murine ATGL and HSL as well as human DAG lipase α, MAGL and the α/β-hydrolase domain 6 ABHD6 (Iglesias et al., 2016). As such, ORlistat has the potential to inhibit the three consecutive steps of TG hydrolysis in murine cells and thus the generation of DAG, MAG and fatty acids. In contrast, ATGListatin is highly specific to murine ATGL (Iglesias et al., 2016; Zechner et al., 2009). The fact that ATGListatin recapitulated the effect of ORlistat on pro-inflammatory cytokines strongly suggests that the first committed step of TG lipolysis is responsible for the observed anti-inflammatory actions in microglia. Further, this suggests that ATGL inhibition is not compensated for by HSL or other lipases as suggested by previous report (Han et al., 2020).

Despite a similar decrease of LPS-induced IL-6, IL-1β and MCP-1 expression by ORlistat and ATGListatin in primary microglia, TNF-α expression remained unchanged. This finding suggests that TG lipolysis regulates the expression of specific pro-inflammatory cytokines in microglia, in agreement with previous studies in co-cultures of macrophages and adipocytes (Xu et al., 2021). However, we cannot rule out that LPS-induced TNF-α expression may be affected at earlier or later time point by ORlistat or ATGListatin as recently shown in primary microglia in response to a 24 h treatment with LPS and ATGListatin (Li et al., 2023). Together, our data demonstrates that inhibition of TG lipolysis downregulates inflammatory responses to LPS in both neonatal and adult microglia in a manner similar to previous observations in neonatal microglia (Li et al., 2023), macrophages and neutrophils (Schlager et al., 2015; van Dierendonck et al., 2022) to suggest that ATGL plays a conserved role in the mitigation of inflammation during acute pro-inflammatory insults in peripheral and brain immune cells.

In agreement with our data *in vitro*, ATGL loss-of-function alleviated LPS-induced IL-6 expression in the hypothalamus and cortex without affecting responses in the hippocampus. MCP-1 expression induced by 1 or to 2 doses of LPS tended to be lower in CX3CR1^ATGL^ KO mice but it was not significant in any of the studied brain regions. Although these results are consistent with our data *in vitro*, they do not fully recapitulate the blunting of pro-inflammatory cytokines observed in primary microglia. This is expected since the samples were from unsorted brain tissue, and thus included all cell types in which LPS promotes pro-inflammatory responses such as endothelial cells and astrocytes. The response from these cells may mask any reduction in responses originating specifically from microglia. Nonetheless, these data suggest that microglial ATGL loss-of-function reduces neuroimmune responses in a manner dependent on the cytokines and brain regions. Importantly, reduced inflammatory responses in CX3CR1^ATGL^ KO mice were associated with changes in behavioural disturbances induced by LPS including sickness- and anxiety-like behaviour. The suppressive effect of LPS on locomotor activity was strongly and consistently reduced in CX3CR1^ATGL^ KO male mice during the EPM and LDB tests. In addition, CX3CR1^ATGL^ KO mice were protected from LPS-induced anxiety as suggested by the lack of response on number of entries and time in the lit zone during the LDB test. Overall, our findings *in vitro* and *in vivo* provide evidence that inhibition or loss of microglial ATGL activity has protective actions on acute neuroimmune and behavioural disturbances induced by LPS. Mitigation of neuroimmune responses by ATGL may extend to other types of brain insults as suggested by a recent study reporting that brain-restricted administration of ATGListatin reduces neural damages induced by stroke in mice (Li et al., 2023)

One potential pathway underlying ATGL actions on microglial LPS responses may implicate reduced release and availability of fatty acids for mitochondrial oxidation. ATGL inhibition reduced FAO in a manner similar to etomoxir thereby demonstrating that basal ATGL-dependent lipolysis in microglia generates substrates for FAO. However, FAO inhibition with etomoxir did not recapitulate the effect of ATGListatin on LPS-induced IL-6 and MCP-1 expression (6 h) and secretion (24 h). These results therefore strongly suggest that the anti-inflammatory actions of ATGL inhibition do not rely on FAO and that FAO does not modulate inflammatory responses to LPS in these experimental conditions. These results are in contrast with a recent study showing that FAO inhibition with etomoxir exacerbates pro-inflammatory responses in microglia activated by myelin debris (Qin et al., 2023). Such difference could be related to different types of inflammatory stimuli (LPS vs. myelin debris) and associated signalling pathways.

Based on our findings, we speculated that ATGL may regulate the generation of fatty-acid-derived inflammatory signals including prostanoids that are arachidonic acid metabolites generated by COX-1 and -2. Using targeted lipidomics, we found that ATGL inhibition led to a dramatic decrease of LPS-induced PGE_2_, PGD_2_ and TXB_2_ secretion and intracellular PGF_2a_ level. While LPS induced the expected changes in COX-1 and -2 expression, co-treatment with ATGListatin did not affect this response, suggesting that ATGL inhibition reduces the amount of arachidonic acid available for prostanoids synthesis. These findings are in agreement with studies reporting that ATGL inhibition or loss-of-function reduces prostanoid synthesis under inflammatory conditions in macrophages (van Dierendonck et al., 2022) and neutrophils (Schlager et al., 2015). Prostanoids, particularly PGE_2_, are well known to activate transcription factors promoting the synthesis of IL-6 and other cytokines (Fiebich et al., 2001; Kawahara et al., 2015; Ricciotti & Fitzgerald, 2011) therefore raising the possibility that mitigation of LPS-induced cytokine responses by ATGL inhibition are secondary to reduced prostanoid signalling.

The broader impact of ATGL inhibition on the microglial lipidome was assessed using untargeted lipidomics. Ceramides were the identified lipid species mostly affected by LPS and/or ATGL inhibition. Interestingly, ATGListatin increased the level of dihydroceramides 18:0/24:0 and 18:0/24:1 while reducing ceramide 18:1/24:0 and 18:1/24:1 levels. LPS treatment showed a similar pattern, though less pronounced, on dihydroceramide and ceramide levels which was exacerbated by ATGL inhibition. ATGListatin also increased sphingomyelin d18:0/18:0 in presence of LPS vs. LPS alone. The *de novo* synthesis of ceramides starts with palmitoyl-CoA which after a series of enzymatic reactions generates dihydroceramides acting as precursor for ceramide synthesis by the enzyme Δ4-dihydroceramide desaturase 1 (DEGS1) (Tzou et al., 2023). Thus, if ATGL activity releases palmitate acting as a precursor for ceramide synthesis, one would expect that ATGL inhibition would reduce the amount of palmitoyl-CoA available and thus the synthesis of dihydroceramides. Given that the opposite is observed on dihydroceramide levels, we could speculate that ATGL inhibition reduces the activity of DEGS1 and the conversion of dihydroceramide into ceramide, resulting in increased dihydroceramide and decreased ceramide levels. It is also possible, and not exclusive, that the reduced ceramide level may result from decreased sphingomyelinase activity, as supported by increased SM 18:0/18:0 level in response to LPS and ATGListatin. Although it is difficult to identify the specific pathway(s) underlying these changes, our findings show for the first time that ATGL regulates ceramide subclasses in microglia including dihydroceramides which are now recognized as bioactive molecules with immunomodulatory properties (Lachkar et al., 2021). The decreased ceramide levels reported here results contrasts with the accumulation of deleterious ceramides reported in ATGL deficient macrophages (Aflaki et al., 2012) and cardiomyocytes (Gao et al., 2015).

Whether or not changes in microglial dihydroceramide and ceramide levels are secondary to decreased prostanoids and cytokines signalling will require additional studies. Overall, the current lipidomic results demonstrate that the anti-inflammatory profile induced by ATGL inhibition in microglia is associated with reduced levels of pro-inflammatory prostanoids and ceramides.

Our findings, in line with recent ones (Li et al., 2023) strongly suggest that ATGL-dependent TG hydrolysis and accumulation of LD in microglia is rather beneficial for neuroinflammation *in vitro* and *in vivo* by alleviating microglial pro-inflammatory responses. These results are in contrast with studies showing that aging, Alzheimer and neurodegenerative diseases or other type of brain insults (e.g. traumatic brain injury) are associated with LD accumulation in microglia (ArbaizarLJRovirosa et al., 2023; Claes et al., 2021; Lee et al., 2023; Marschallinger et al., 2020; Zambusi et al., 2022). This difference may be related to the duration or nature of insults. It is conceivable that acute pro-inflammatory insults increase LD number which mitigates inflammation but that chronic accumulation of LD in microglia promotes or aggravates neuroinflammation.

Collectively, our findings identify ATGL-dependent LD lipolysis as a key regulator of microglia reactivity and neuroinflammation during acute inflammatory insults that may have implications for our understanding of the pathways regulating microglia activity in physiological and pathophysiological states. Further studies will be required to elucidate the role of microglial ATGL and LD lipolysis in CNS pathologies associated with neuroinflammation and microglial activation such as mood disorder, obesity and Alzheimer.

## Declarations

### AUTHORS CONTRIBUTIONS

J.L.R., A.I.M.P, F.B., R.M., D.M., D.R, K.B., A.C. and C.L. helped with colony management and genotyping, cell culture, qPCR, ELISA, fatty acid oxidation, LD imaging, FACS experiments and behaviour tests. M.R., A.F. and C.D. carried out LC-MS/MS for untargeted lipidomics, data analysis and interpretation, and contributed to manuscript preparation. K.G. and J.-F.B. carried out LC-MS/MS for targeted lipidomics, data analysis and contributed to manuscript preparation. J.L.R., A.I.M.P., F.B., D.M., K.B., X. F., C.M.-D., S.L., N.A., S.F. and T.A. contributed to conceptualization, experimental design, data interpretation and manuscript revisions. J.L.R. and T.A. wrote the manuscript.

## ACKNOWLEDGEMENTS

We are grateful to Dr Grant Mitchell for the ATGL-floxed mice. We thank the CRCHUM cytometry core facility for their help with FACS studies. This work was supported by grants from the National Natural Sciences and Engineering Research Council (NSERC, RGPIN/04798), the Canadian Institutes of Health Research (CIHR, PJT153035) and Réseau de recherche en santé cardiométabolique, diabète & obésité from Fonds de Recherche Québec-Santé (CMDO-FRQS) to T.A., and grants from Fondation pour la Recherche Médicale (FRM), Fédération pour la Recherche sur le Cerveau (FRC) and Institut Benjamin Delessert to X.F.. A.C. was supported by a fellowship from INRAE/Bordeaux INP. S.F., M.R. and T.A. were supported by a salary award from FRQS. J.L.R., R.M and D.M. were supported by doctoral and postdoctoral FRQS fellowships and PhD fellowships from the Neuroscience Department. A.I.M.P was supported by a fellowship from Montreal Diabetes Research Center and Faculty of Medicine. C.L. obtained postdoctoral awards from Fondation d’Aide pour la Recherche sur la Sclérose en Plaques (ARSEP), FRQS and CIHR.

## ETHICS APRROVAL

All animal studies were conducted in accordance with guidelines of the Canadian Council on Animal Care (protocol #CM19018TAs) and in accordance with the European directive 2010/63/UE and approved by the French Ministry of Research and local ethics committees (APAFIS#: 33951).

## DISCLOSURES

Authors have no conflict of interest to declare.

## DATA AVAILABILITY STATEMENT

Data are available on reasonable request from the authors.

## Notes

### Competing Interest Statement

The authors have declared no competing interest.

