## Supplemental figures and legends for "Microglial adipose triglyceride lipase regulates neuroinflammatory and behavioural responses to LPS"

**Supplementary figure 1**

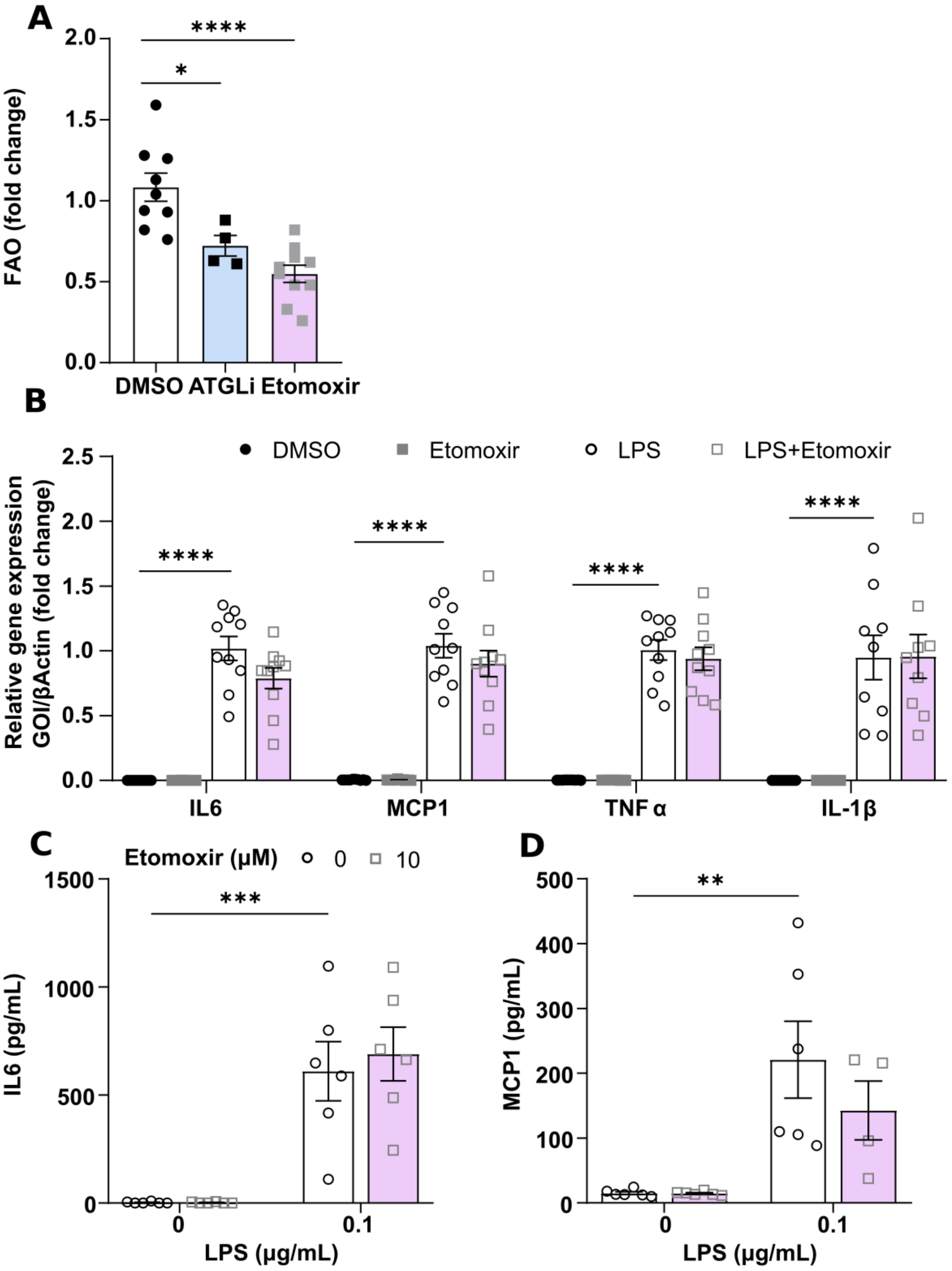

***Supplementary figure 1 – Inhibition of mitochondrial fatty acid oxidation does not alter LPS-induced cytokine expression and secretion.***

Effect of 0.1 µg/mL LPS ± etomoxir (10 µM, CPT1a inhibitor) on cytokine expression and secretion. **A.** Oleate oxidation after 3 h treatment with 50 µM ATGListatin or 10 µM etomoxir (n=4-10). *$p < 0.05$; **** $p < 0.0001$; One-way ANOVA with post-hoc Tukey's. **B.** Relative gene expression of cytokines after 6 h treatment with LPS ± etomoxir (n=9-10). Extracellular cytokine concentration of **C.** IL-6 (n=6) and **D.** MCP-1 (n=4-6) 24 h post-treatment. ** $p<0.01$; *** $p<0.001$; **** $p<0.0001$; Two-way ANOVA with post-hoc Tukey.

**Supplementary figure 2**

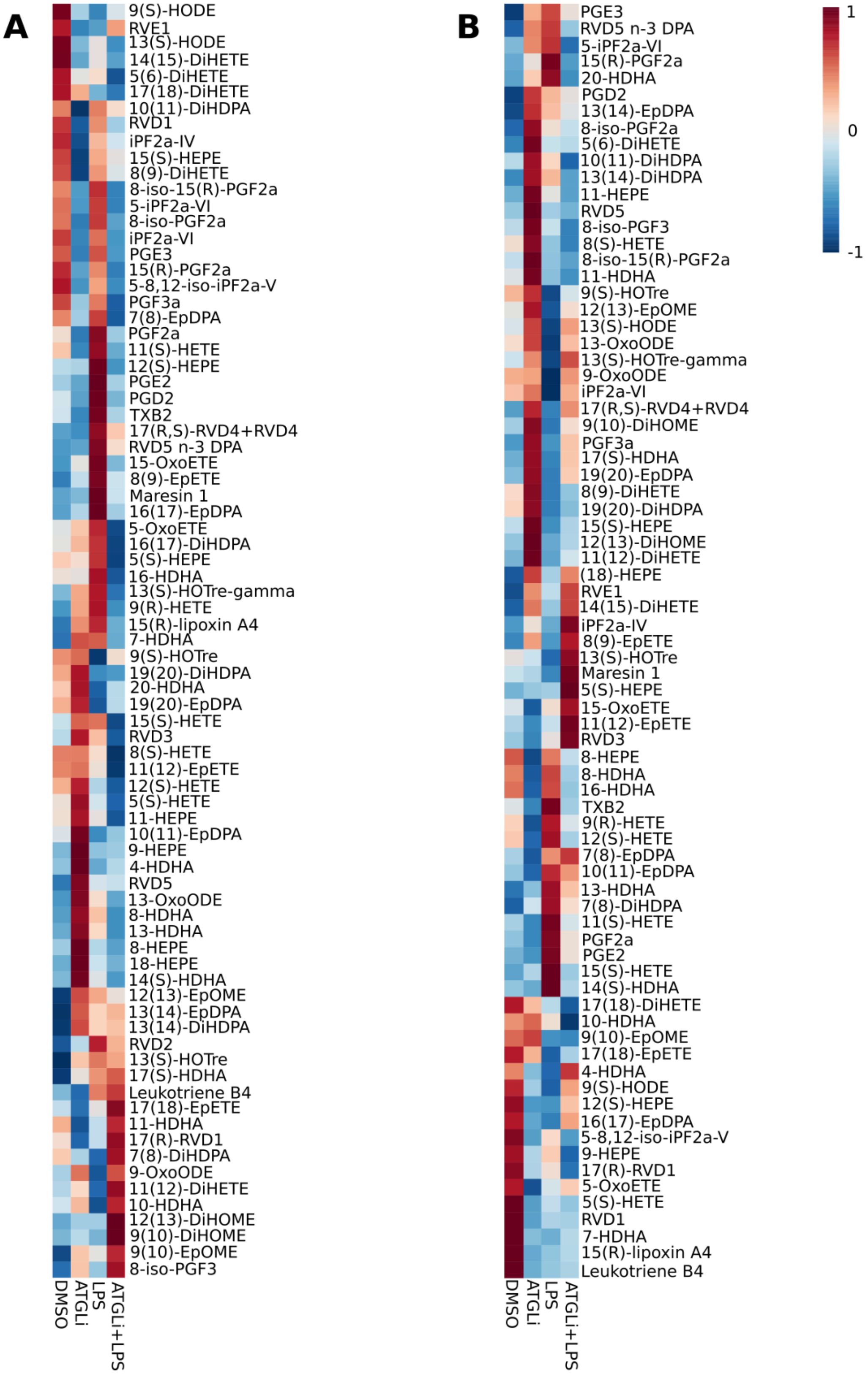

***Supplementary figure 2 – Inhibition of ATGL alters LPS-induced changes in detected oxylipins.***

Effect of 24 h 0.1 µg/mL LPS ± 50 µM ATGListatin on **A.** extracellular and **B.** intracellular concentrations of oxylipins as detected by LC-MS/MS (n=6).

**Supplementary figure 3**

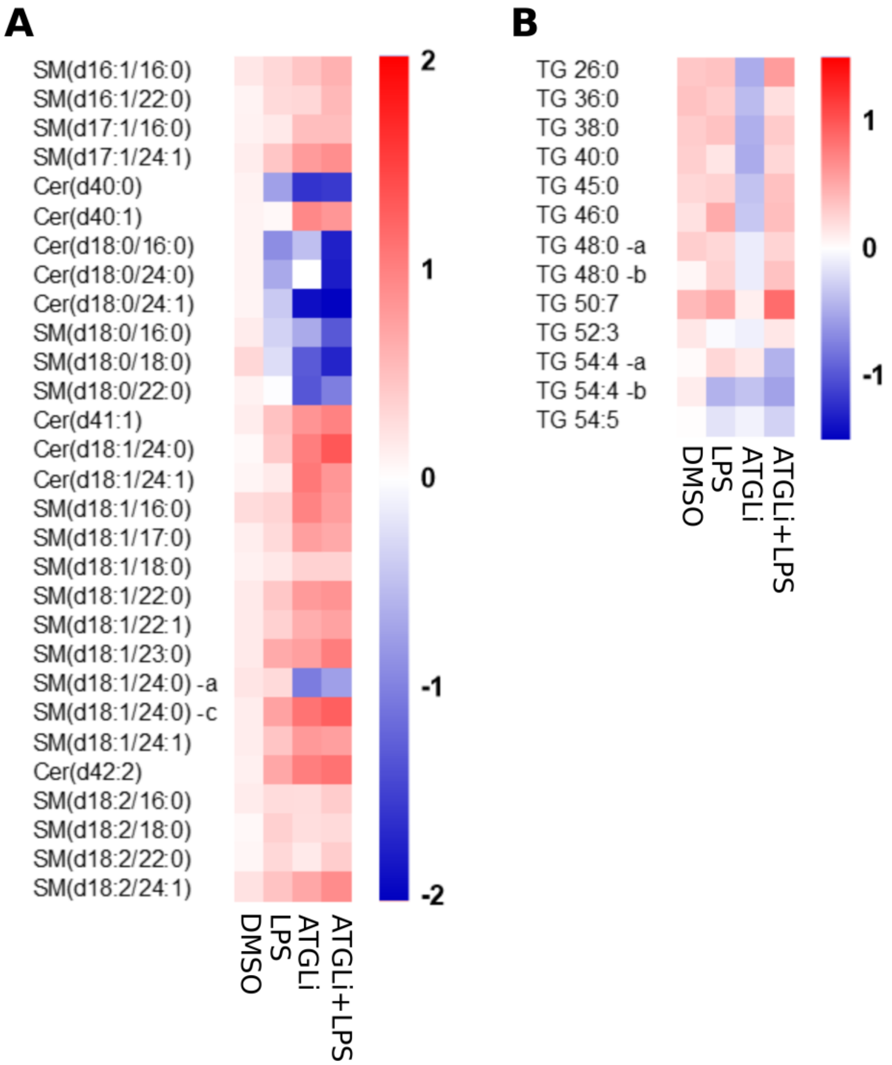


***Supplementary figure 3 – Inhibition of ATGL alters accumulation of sphingolipids and triglycerides***

Effect of 24 h 0.1 µg/mL LPS ± 50 µM ATGListatin on intracellular lipid species. **A.** Heatmap of changes in identified ceramides and sphingomyelin. **B.** Heatmap of fold changes in identified triglycerides (n=6).

**Supplementary figure 4**

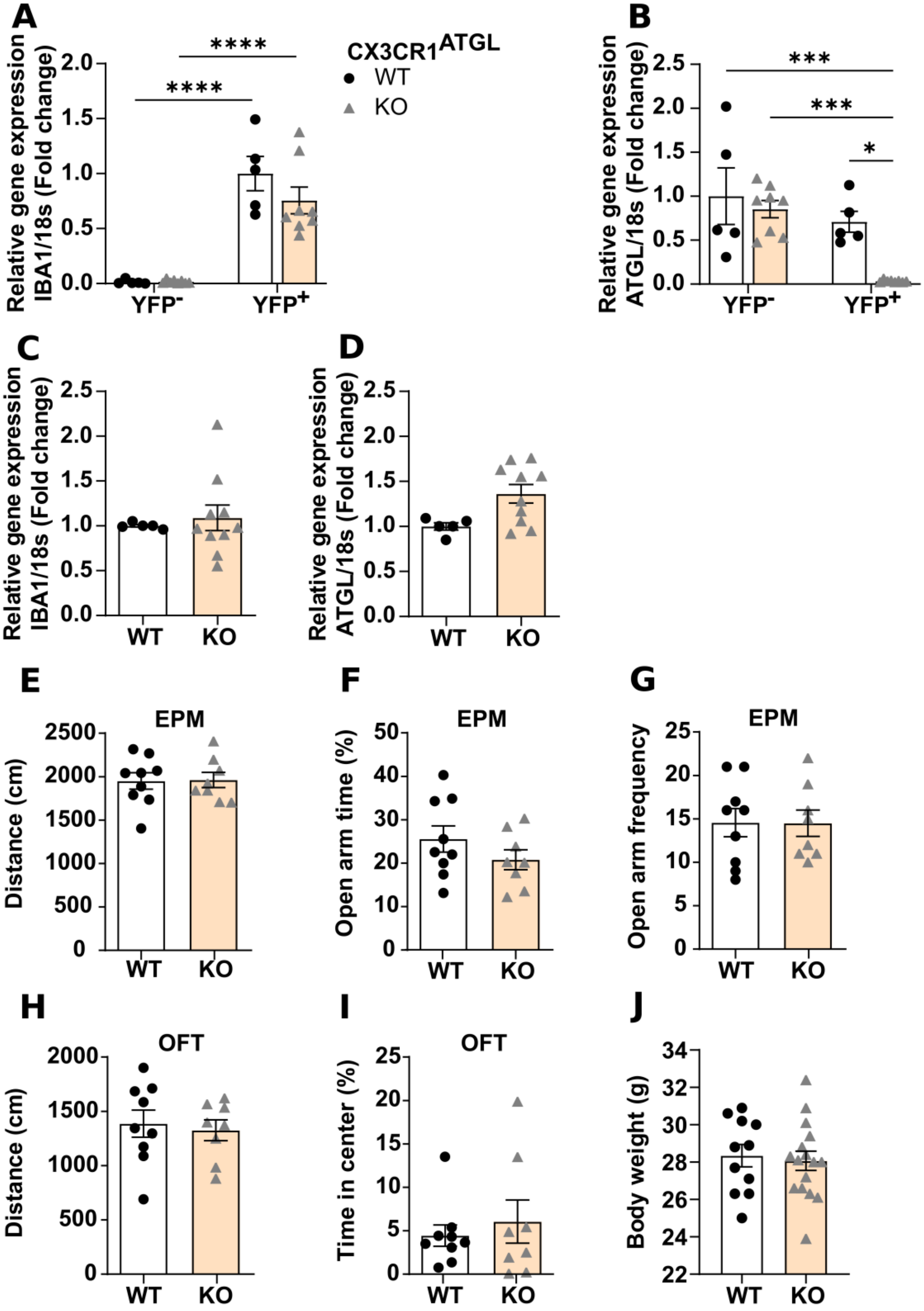

***Supplementary figure 4 – Validation and anxiety-like behaviour in mice with a specific and inducible loss of ATGL from microglia.***

Gene expression of **A.** IBA-1 and **B.** ATGL in FACS sorted $CD11b^{+}/YFP^{+}$ (microglia) and $CD11b^{-}/YFP^{-}$(non-microglia) cells isolated from the brain of $CX3CR1^{ATGL}$WT and KO animals (n=5-8). * $p<0.05$, *** $p<0.001$, **** $p<0.0001$, two-way ANOVA with post-hoc Tukey. Gene expression of **C.** IBA-1 and **D.** ATGL in FACS sorted $CD11b^{+}$ (macrophage) cells isolated from the spleen of $CX3CR1^{ATGL}$WT and KO animals (n=5-8). Elevated plus maze (EPM): **E.** Distance moved, **F.** percentage time in open arm and **G.** frequency of entries to open arm (n=8-9). Open field test (OFT): **H.** distance moved **I.** percentage of time spent in centre (n=8-9). **J.** Body weight (n=11-16). Student's t-test.
